# Single-array measurements reveal non-uniform, mosaic-like chemosensory arrays in bacteria

**DOI:** 10.1101/2025.10.01.679710

**Authors:** Vered Frank, Nir Livne, Moriah Koler, Ady Vaknin

**Affiliations:** The Racah Institute of Physics, The Hebrew University, Jerusalem 91904, Israel

**Keywords:** chemotaxis, cell signaling, receptor array signaling, signal integration

## Abstract

Motile bacteria use supramolecular arrays to detect effector gradients in their environment. In *Escherichia coli*, thousands of chemoreceptor molecules with diverse sensory properties cooperatively modulate the kinase activity of these arrays and, via phosphotransfer to a diffusible response regulator, control the cell’s swimming behavior. Various methods have been used to study these sensory arrays in live cells, from population-level assays to single-cell measurements, revealing hierarchical coupling interactions that underlie their remarkable sensory properties. However, measuring the responses of individual arrays has remained a challenge. Here, by combining the kinase and response regulator into a functional hybrid protein that resides within the array, we directly measured the kinase responses of individual arrays in live cells. These measurements revealed highly diverse and growth-phase-dependent sensory properties of individual arrays. Even arrays within the same cell were not substantially correlated. We also observed dynamic shifts in receptor occupancy within individual arrays. Overall, these data suggest that each array contains a ‘frozen’ non-uniformity, reflecting its unique assembly history and resulting in a mosaic arrangement of cooperative signaling regions, each with distinct receptor content. Consistent with this view, measured dose-responses of individual arrays mostly exhibit low cooperativity. Interestingly, signal integration in such non-uniform arrays is expected to inherently vary with the size of the cooperative regions.

## Introduction

The ability of motile bacteria to swim toward beneficial effectors (attractants) and away from harmful effectors (repellents)^1^ influences many aspects of bacterial ecology^2^ and human health^3, 4^. This behavior, termed chemotaxis, relies on sensory complexes comprising thousands of transmembrane receptor molecules that detect effector gradients along the swimming trajectory of the cell and modulate its swimming behavior accordingly. The architecture of these extended sensory arrays in *E. coli* is illustrated in Fig. 1A^5-8^. Each cell can have several arrays that preferentially nucleate at the cell poles and expand over multiple cell divisions^9-11^. The most abundant chemoreceptors are Tar and Tsr, which primarily detect aspartate and serine, respectively^12, 13^. The minimal core signaling unit consists of two receptor trimers associated with a homodimeric kinase (CheA) and two adaptor proteins (CheW)^14-17^. These core complexes further aggregate through direct contacts between the kinase CheA.P5 domain in one complex and the CheW protein in another, forming a networked CheA–CheW baseplate and extended arrays comprising thousands of receptor molecules. Although receptor-mediated signals can be integrated within core complexes^17, 18^, extended arrays exhibit enhanced cooperativity due to long-range allosteric interactions^8, 16^, which lead to enhanced sensitivity and increased dynamic range^19-21^.

**Figure 1.**
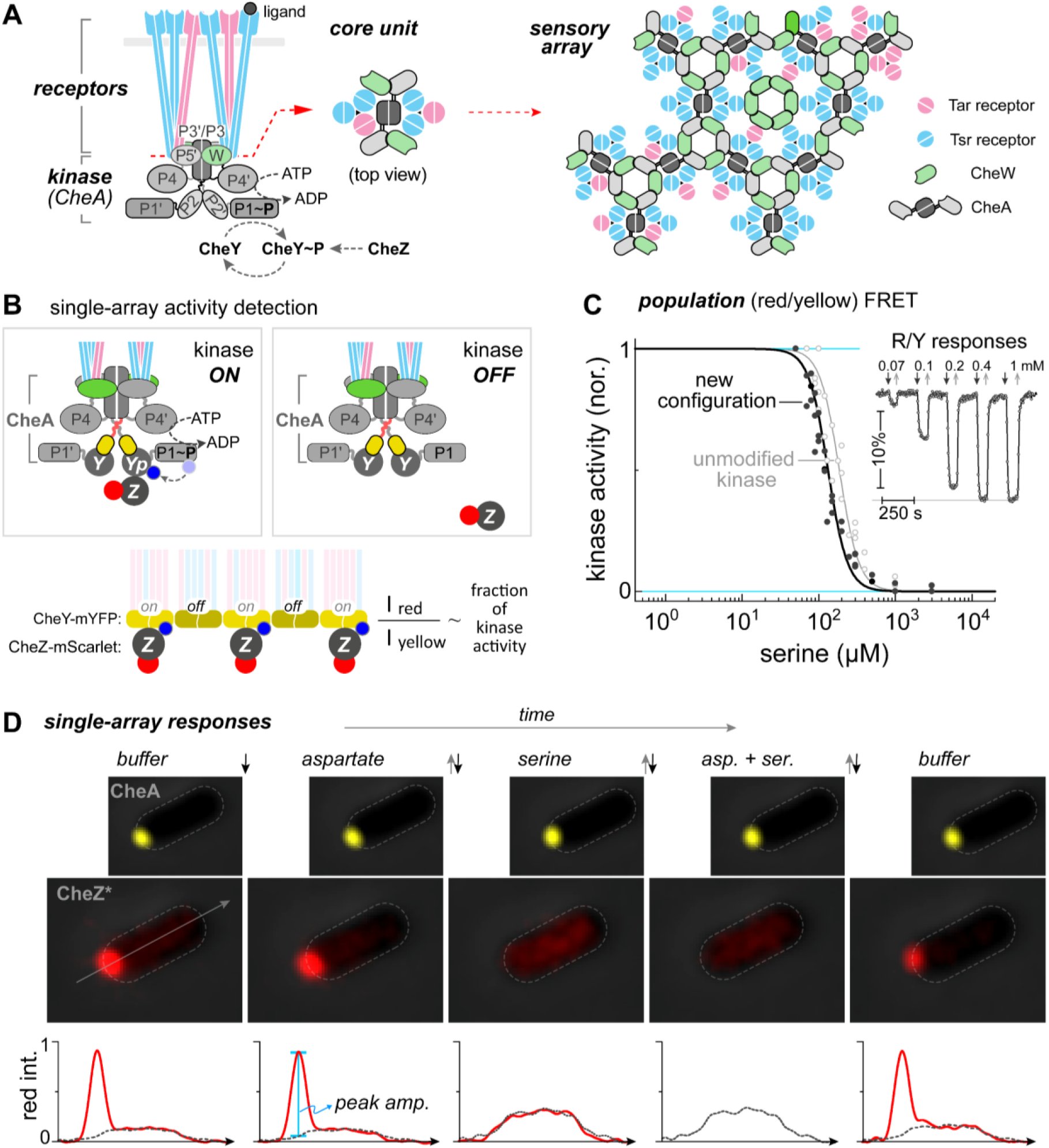
A method for measuring kinase responses in individual receptor arrays. (**A**) Schematic of the core signaling complex, composed of a kinase dimer (CheA), two adaptor proteins (CheW), and six membrane-bound receptors arranged as two heterotrimers of homodimers. Also shown are the reactions involving the response regulator CheY: its phosphorylation by the kinase CheA and dephosphorylation by the phosphatase CheZ (left panel). The organization of these complexes in the extended array is shown in the right panel. (**B**) Schematic of the hybrid kinase/response-regulator protein (CheA::CheY– mYFP), in which CheY–mYFP replaces the P2 domain of CheA. Kinase activity is reported through the activity-dependent association of fluorescently tagged CheZ(F98S) with phosphorylated CheY during catalysis. (**C**) Serine dose-dependent kinase inhibition in VF17 cells was measured by population FRET between CheA::CheY–mYFP and CheZ(F98S)–Scarlet (black symbols). For comparison, corresponding measurements are shown for cells with native CheA (UU2828, *ΔcheYcheZ*) expressing tagged CheY and CheZ from a plasmid (gray symbols; pAV109). *Inset*: Representative serine-induced responses in VF17 cells. (**D**) Time-series fluorescence images of a single VF17 cell from an exponential-phase culture (OD 0.45) exposed to different stimuli (3 mM; as labeled), recorded using either ‘yellow’ (upper row) or ‘red’ (lower row) filter sets. Below each image, the corresponding red fluorescence intensity profile along the cell axis (arrow in the leftmost image) is plotted, corrected for photobleaching (Fig. S3). For reference, the intensity profile for the serine+aspartate condition is overlaid in all other conditions (dashed lines), normalized to the cell-body intensity (where no arrays are present). Peak amplitude (blue bar) was calculated by subtracting background fluorescence at the array location due to unbound CheZ.

Stimuli detected by the receptors modulate the kinase activity of the array, which in turn controls the phosphorylation state of the cytoplasmic response regulator CheY. In its phosphorylated form, CheY binds to the flagellar motor and promotes random changes in swimming direction. In steady state, these reactions are balanced by a dedicated phosphatase (CheZ), which removes phosphate groups from CheY to ensure rapid sensory responses. Utilizing this steady-state balance, kinase activity can be monitored in live cells by measuring FRET between fluorescently tagged CheY and CheZ^20, 22^. Such FRET-based assays have been used to measure kinase responses averaged across entire cell populations, revealing key properties of these sensory complexes, including cooperative responses at multiple levels that enhance receptor detection sensitivity and dynamic range^16, 18-20, 23-25^. These measurements have also contributed to a more quantitative understanding of sensory-array signaling^25-28^. Kinase responses have also been detected in single cells^29-32^, revealing cell-to-cell variability and temporal fluctuations. However, measuring the kinase activity of individual arrays within live cells has remained a challenge.

Here, we present an assay that enables the direct measurement of kinase responses in individual sensory arrays within live bacterial cells. This assay relies on combining the kinase CheA with the response regulator CheY to create a fluorescently tagged hybrid protein (CheA::CheY–mYFP), which permanently localizes to the array. In this configuration, the affinity of CheZ for CheYp in each array provides a direct readout of its kinase activity. In addition, by also tagging either Tar or Tsr in distinct strains, we could measure their occupancy relative to CheA within individual arrays. The observations presented here suggest that receptor arrays are highly non-uniform, with a mosaic arrangement of cooperative subregions, each with distinct receptor content. This non-uniformity likely reflects the unique assembly history of each array as it develops over multiple generations. Interestingly, internal non-uniformity in array composition also implies that the size of the cooperative sub-regions, which determines the scale over which this non-uniformity is averaged, can fundamentally influence how signals are integrated within the array.

## Results

### Measuring kinase activity in individual arrays

As illustrated in Fig. 1A (left), ligand binding to the chemoreceptors modulates the phosphorylation rate of the CheA.P1 domain —referred to here as ‘kinase activity’— mediated by interaction of this domain with the catalytic domain complex CheA.P3-4. The diffusible response regulator CheY acquires phosphoryl groups from CheA.P1, forming CheYp^33^. CheYp then diffuses from the array to the cell’s flagellar motors to trigger changes in swimming behavior. These reactions are balanced by CheZ-mediated dephosphorylation of CheYp.

To follow kinase activity at the array, we made a strain (VF17) having two chromosomal modifications (Fig. 1B and *Materials and Methods*): First, we replaced the CheA.P2 domain with fluorescently tagged CheY, yielding a hybrid kinase–phosphotransfer protein CheA::CheY– mYFP. This design ensures that phosphotransfer from CheA.P1 to CheY occurs internally within the array, while CheYp remains part of the array. Second, we fused the phosphatase CheZ to mScarlet and introduced a point mutation (F98S) that abolishes direct interaction between CheZ and CheA, while preserving its phosphatase activity toward CheYp^29, 34^. Thus, in this new configuration, the tagged CheZ associates with the array only through transient interactions with CheYp during catalysis (Fig. 1B). As a result, the ratio of red (CheZ–mScarlet) to yellow (CheA::CheY–mYFP) fluorescence in each array directly reflects the rate of CheYp dephosphorylation, given by [CheYp·CheZ]·K_cat_, normalized by the size (CheA content) of the array. Thus, at steady state, this red-to-yellow ratio provides a readout of the kinase activity per CheA molecule in the array:

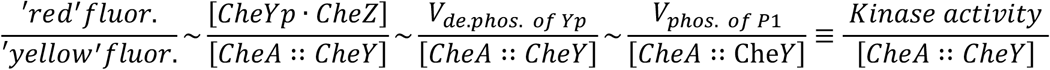

where *V*_*de*.*phos of Yp*_ and *V*_*phos. of P1*_ are the velocities of the dephosphorylation and kinase reactions. Notably, these relationships hold for each array, independently of other arrays that may be present in the cell.

Some inherent cross-talk between arrays within the same cell might be expected due to shared CheZ (*Supplementary Text 1*). To estimate this cross-talk, we first evaluated the fraction of CheZ proteins bound to arrays in the unstimulated state (Fig. S1A). We found that the amount of CheZ bound to the arrays was up to approximately 40%, and a few cells containing more than one array, in which this fraction exceeded 50%, were excluded from further analysis. Based on these data, we estimated the mutual influence of kinase activity between different arrays in the same cell to be about 10%, rising to about 15% in extreme cases where a very small array coexists with a much larger one (Fig. S1).

To verify that the hybrid protein retains its dual function, we measured FRET between CheA::CheY–mYFP and CheZ–mScarlet in cell populations of the new VF17 strain. Clear responses to an attractant ligand (serine) were indeed observed (Fig. 1C, inset), indicating that CheZ association with the array is ligand dependent. This also implies that sensory inputs modulate both CheA autophosphorylation and phosphotransfer to CheY. Because these cells lack the adaptation enzymes, the responses persisted until the ligand was removed. Moreover, the ligand dose-dependent kinase responses closely matched those of native arrays with unmodified kinase (Figs. 1C and S2), exhibiting similar Hill coefficients (slopes) and ligand sensitivities (K_1/2_). These results are also consistent with previous measurements of mixed arrays^8, 23^. The minor shift in K_1/2_ observed in Fig. 1C likely reflects slight difference in receptor expression (Fig. S2), either due to the added label in the new VF17 strain or due to modifications in the reference strain, namely, the *cheYcheZ* deletions and/or the plasmid-borne expression and associated antibiotic requirements, which also affect growth rate. Finally, we confirmed that the arrays remained stable and unaffected by ligand binding (Fig. 1D, yellow). We therefore conclude that the hybrid protein does not significantly alter the sensory responses of the arrays.

An example of how ligand binding to the receptors modulates the association of CheZ with an individual receptor array is shown in Fig. 1D (lower images, red). The corresponding intensity profiles of CheZ–mScarlet fluorescence across the cell are shown below (red lines), corrected for photobleaching (see *Materials and Methods* and Fig. S3). Upon addition of serine (3 mM), CheZ dissociated from the array, indicating that the kinase activity within the array had switched off. Interestingly, despite the potential presence of all five receptor types in this strain, in the specific array shown in Fig. 1D, aspartate addition (3 mM) had no observable effect on CheZ association, implying unchanged kinase activity. Finally, in the presence of both serine and aspartate, CheZ consistently exhibited no significant affinity for the array, representing the kinase-OFF state.

To quantify CheZ association with the array, we used the peak (red) fluorescence intensity in the array and subtracted the background fluorescence at the array location due to unbound CheZ (Fig. 1D, lower plots, dashed lines), which, in turn, was evaluated based on the distribution of CheZ in the presence of both serine and aspartate (kinase-OFF state). To account for changes in the total unbound CheZ in the cell upon stimulation, the red intensity at the cell body—where no array is apparent— in the presence of both serine and aspartate was normalized to the cell body intensity in each case (Fig. 1D). While Fig. 1D captures the essence of the analysis used here to evaluate the ligand responses, the actual analysis was mostly carried out in two dimensions, as described in *Supplementary Methods*. Additional examples are shown in Fig. S4.

### Distributions of single-array responses

As expected, the kinase activity in the unstimulated state—indicated by the amount of CheZ–mScarlet associated with the array—scales approximately linearly with the array size, indicated by the CheA::CheY–mYFP fluorescence (Fig. 2A, left). Interestingly, however, the efficiency of kinase activation, namely, the kinase output per CheA (Fig. 2A, right), is somewhat higher in arrays from cells at the early or late stationary growth phase (OD 1.7 or 3.5), compared with cells in the exponential phase (OD 0.15 or 0.45). The growth curve of these cells is shown in Fig. 2A (right plot, inset), and the number of arrays per cell is shown in Fig. S5, generally consistent with the observations in ref. 11.

**Figure 2.**
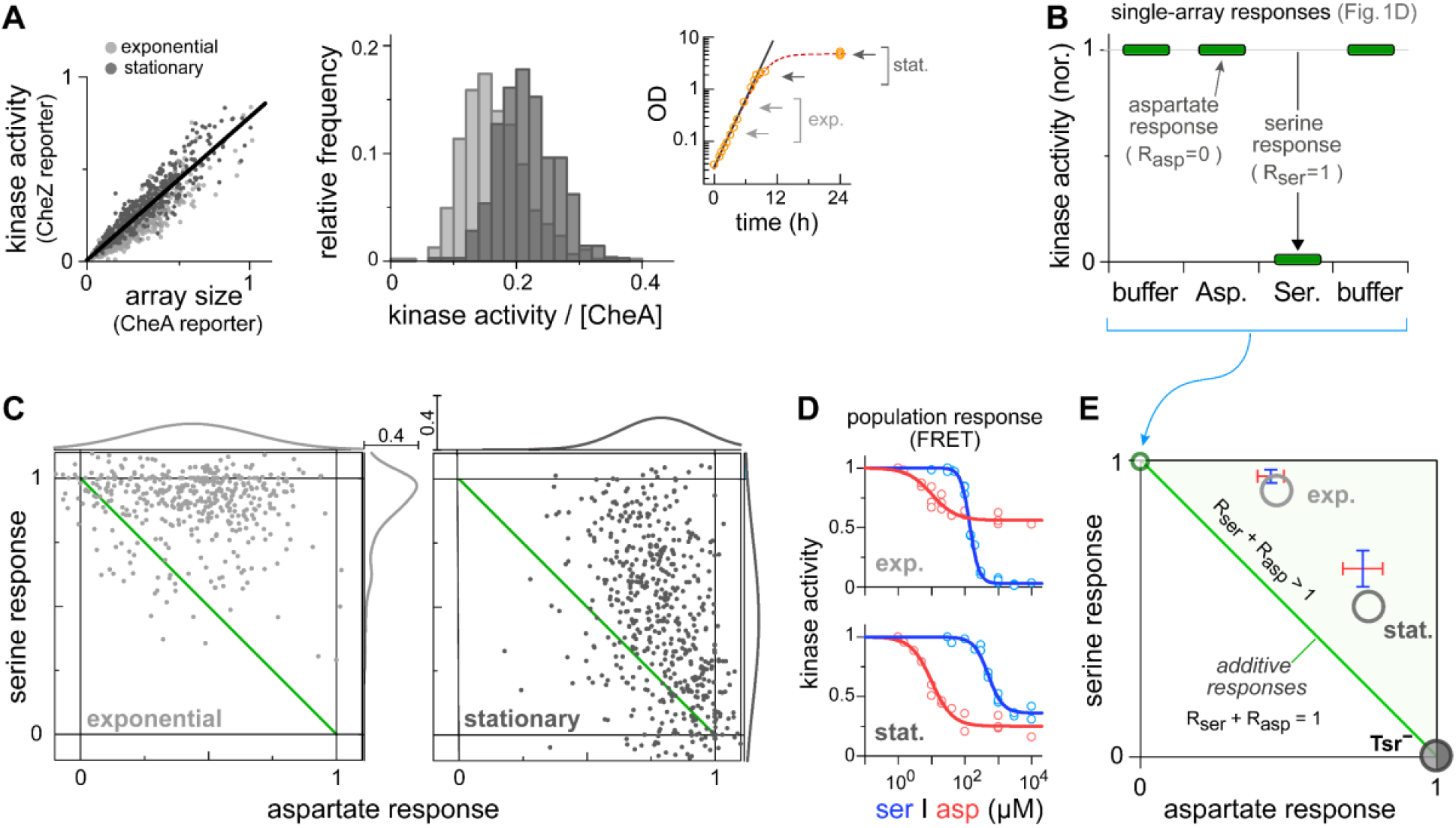
Signaling properties of individual arrays. (**A**) *Left panel*: Total kinase activity in unstimulated arrays (CheZ–mScarlet fluorescence, divided by 800) plotted against the amount of kinase molecules in each array (CheA::CheY–mYFP fluorescence, divided by 3500), for VF17 cells in exponential (light gray) or stationary (dark gray) growth phases. *Right panel*: Distributions of kinase activity per CheA molecule in the arrays. Arbitrary units are used in both plots. *Inset:* Growth curve of VF17 cells. (**B**) Quantified responses of the array shown in Fig. 1D, normalized to the unperturbed kinase activity measured in buffer, representing the fraction of kinase activity inhibited by each stimulus. (**C**) Responses of individual arrays, such as those shown in part (B) and Fig. S4, mapped onto a ‘Ser–Asp’ response plot, in which the fraction of kinase activity inhibited by serine is plotted against that inhibited by aspartate. Shown are responses of individual arrays in exponential (OD 0.15 and 0.45; 480 arrays) and stationary (OD 1.7 and 3.5; 500 arrays) growth phases to either serine (3 mM) or aspartate (3 mM). Also shown are the response distributions (relative frequency) along the axis. (**D**) Population-level FRET dose-response behavior of VF17 cells in the exponential (OD 0.45; upper plot) and stationary (OD 3.5; lower plot) growth phase to either serine (blue) or aspartate (red). Lines represent fit to Hill function with K_1/2_ and Hill coefficient being, respectively: exponential phase – 140 μM, 3.2 (serine) and 10 μM, 1.4 (aspartate); stationary phase – 500 μM, 2 (serine) and 10 μM, 1.4 (aspartate). (**E**) Summary plot: the Ser–Asp plot is shown with the ‘additivity’ line marked by a green line. Also indicated are the position of the array shown in (B) (upper-left corner); the weighted average of the responses shown in (C) (gray circles); the responses extracted from the population FRET data in (D) (blue-red bars); and the response exhibited by cells lacking Tsr receptors (black circle, Fig. S6).

The responses of the array shown in Fig. 1D are plotted in Fig. 2B, normalized to the unperturbed kinase activity state, indicated by the red fluorescence peak in the absence of any stimulus. These normalized responses represent the fraction of kinase activity in the array inhibited by each stimulus, and notably, they reveal the relation between the effects of these stimuli on single arrays (see Fig. S4 for additional examples). To compare different arrays, the responses of each array were represented by a single point in a ‘Ser–Asp’ response plot (Fig. 2C), where the fraction of kinase activity responsive to serine is plotted against that responsive to aspartate. These responses were measured in cell populations at either the exponential or stationary growth phase. Individual arrays exhibited a wide range of sensory properties with a semi-continuous distribution. Additionally, consistent with previous observations^35^, the repertoire of these sensory properties shifted from serine-biased in the exponential phase to aspartate-biased in the stationary phase. Notably, however, the orthogonal ligand in each case (aspartate and serine, respectively) still elicited a wide range of responses. The corresponding population-level (FRET) dose-response measurements of these cells also showed a similar bias shift (Fig. 2D). This shift was consistent with the weighted-average values obtained from the single-array responses (Fig. 2E), in which the contribution of each array was weighted by its peak mYFP fluorescence, used as a proxy for array size, and thus for the kinase contribution of that array. A slight shift is apparent in the stationary phase, which could result from this approximation.

Three regions in the Asp–Ser plot are notable (Fig. 2E): First, the region above the diagonal (green background), where the sum of the responses to serine and aspartate is larger than the response to both stimuli combined, implying that a fraction of the kinase molecules in the array can be inhibited by either Tar or Tsr. Second, the region along the diagonal (green line), where the sum of the individual responses equals the response to both stimuli combined. Such ‘additive’ behavior may occur if Tsr and Tar each regulate distinct subsets of kinase molecules. Third, the region below the diagonal, where the sum of the individual responses is smaller than the response to both stimuli combined, implying that some fraction of the kinase molecules can only be inhibited when both stimuli are present. Clearly, most of the single-array responses observed in Fig. 2C fall above the additivity line^36^, indicating robust overlap in the kinase activity controlled by Tar and Tsr within each array.

As expected, when the experiments were repeated with cells lacking the Tsr receptor, a complete aspartate bias was observed in both growth phases (Figs. S6 and 2E, black circle). This result is also consistent with the Aer, Tap, and Trg receptors having only a minor influence under the conditions tested here, in line with their typically low abundance, constituting only a few percent of the total receptor population^37^. These receptors however may play a more significant role in the subpopulation of arrays that fall below the ‘additivity’ line in Fig. 2C. Additionally, response patterns similar to those observed in Fig. 2C were also observed for a subpopulation of arrays that present alone in their respective cells (Fig. S7), further supporting the conclusion that the potential CheZ-mediated interference between arrays does not significantly alter the results shown here (see also *Supplementary Text 1*).

The pairwise correlations of different arrays within the same cell are shown in Fig. 3 (upper plots), where the response of one array is plotted against the corresponding response of another. In each case, the responses are shifted by the mean and scaled by the standard deviation of the respective dataset. Also shown in Fig. 3 (lower plots, gray bars) are the distributions of the pairwise differences, 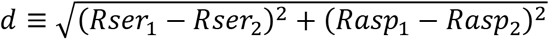, where *Rser*_1,2_ and *Rasp*_1,2_ represent the responses of individual arrays (1 and 2). For comparison, the corresponding distributions obtained by randomly pairing arrays from the respective datasets, are also shown (lower plots, green bars), mostly overlapping the distribution of the same-cell pairs.

**Figure 3.**
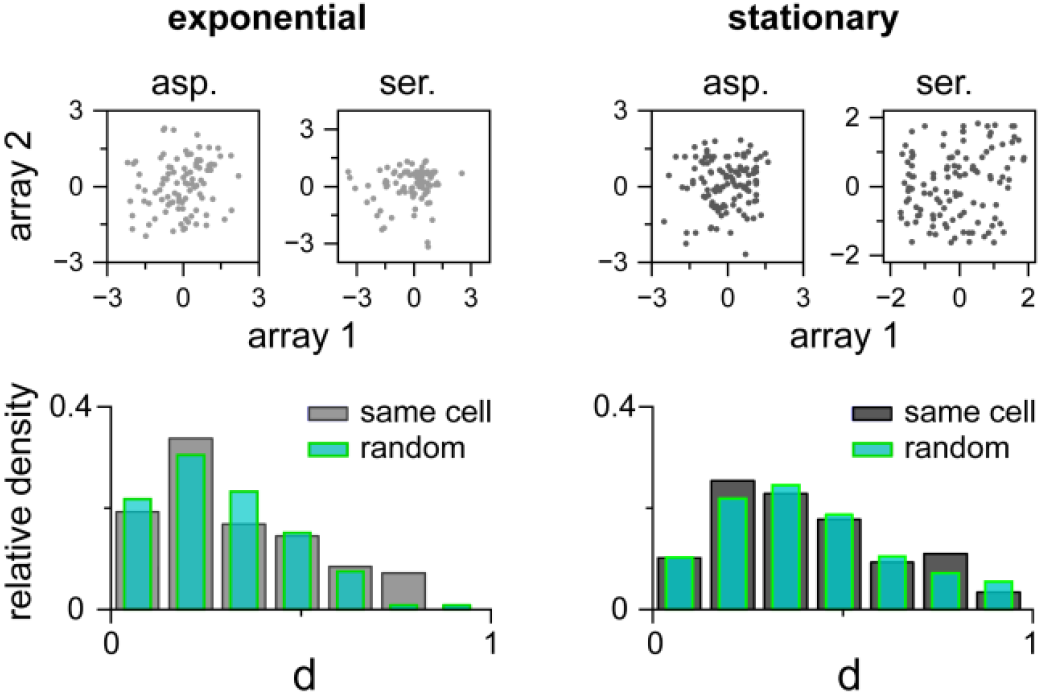
Responses of different arrays within the same cell are not strongly correlated. *Upper plots:* The response of one array to a given ligand is plotted against the response of another array within the same cell to the same ligand. To account for the overall shift in the response distribution, each value was shifted by the mean and normalized by the standard deviation of the respective dataset. Data are shown for responses to serine and aspartate in cells from both the exponential (left) and stationary (right) growth phases (∼100 arrays in each case). The corresponding Spearman correlation values (left to right) are: 0.2 (p=0.04), 0.13 (p=0.18), 0.04 (p=0.65), and 0.13 (p=0.12). *Lower plots:* Distributions of the pairwise 2D distances (d, see text) between responses of arrays within the same cell. Data are shown for actual array pairs (gray bars) and for randomly paired arrays from the general population (green bars).

Overall, these data reveal diverse and growth-phase-dependent sensory properties across chemoreceptor arrays, with arrays in the same cell not significantly more correlated than those in the general population.

### Receptor content and assembly of single arrays

To monitor the fractional occupancy of each array by Tar or Tsr, we constructed strains analogous to VF17, carrying CheA::CheY– mYFP, and either Tar or Tsr tagged with mScarlet, each in a separate strain (tagged CheZ was absent in these strains). In this setup, the ratio of red (mScarlet, receptors) to yellow (mYFP, CheA) fluorescence in each array reflects the amount of Tar or Tsr relative to the size of the array baseplate (Fig. 4). By tagging only one receptor type in each strain, we reduced the total number of tagged receptors, thereby minimizing potential steric effects.

**Figure 4.**
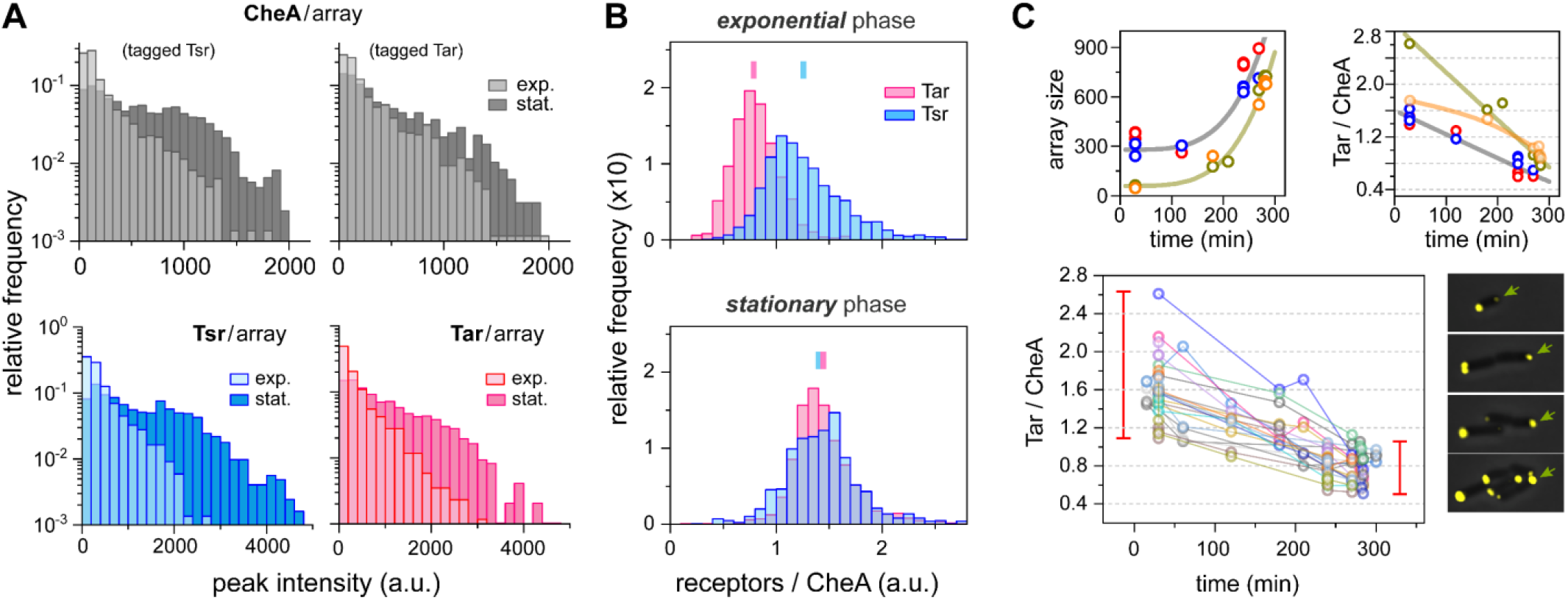
Tar and Tsr occupancy in arrays. Receptor occupancy, defined as the receptor-to-CheA ratio, reflects the fraction of potential array positions occupied by either Tar or Tsr receptors. To assess occupancy, we used cells in which either Tsr (strain MK27) or Tar (strain MK30) was tagged with mScarlet, in addition to CheA::CheY–mYFP. (**A**) *Upper plots:* Distributions of array sizes (mYFP fluorescence) for both strains in exponential and stationary growth phases, as indicated. *Lower plots:* Corresponding distributions of receptor levels in each array (mScarlet fluorescence). (**B**) Distributions of the mScarlet-to-mYFP fluorescence intensity ratio in individual arrays, representing either Tsr-to-CheA (blue) or Tar-to-CheA (red) ratios. Data are shown for cells in exponential (upper plot) and stationary (lower plot) growth phases. Means are indicated by bars. (**C**) *Upper plots:* Growth of individual arrays (left; mYFP fluorescence) and the corresponding changes in Tar occupancy within those arrays (right; red/yellow fluorescence) following a shift from stationary to exponential phase, achieved by placing stationary-phase cells onto an agarose hydrogel pad with fresh growth medium. Sample images are shown (lower-right panel), with the arrow highlighting the same array over time. *Lower plot*: Additional traces of Tar occupancy in individual arrays, showing an overall shift (red bars) consistent with the population-level data in (B). Different symbol colors correspond to different arrays. Fluorescence intensity was corrected for bleaching (see Fig. S3).

We observed that the size of the arrays and, correspondingly, the amount of Tar or Tsr tended to increase as cells approached stationary phase (Fig. 4A). Moreover, in the exponential phase, the Tar fraction in the arrays (Tar/CheA) tended to be lower than that of Tsr, whereas in the stationary phase the receptor distributions became similar (Fig. 4B). This shift in receptor occupation is consistent with previous observations^35, 38^, albeit more moderate^35, 38^, possibly due to different growth conditions (see *Materials and Methods*). Interestingly, the total receptors (Tar + Tsr) to CheA ratio also increased in the stationary phase (Fig. 4B), potentially accounting for the enhanced kinase activity observed in Fig. 2A.

To follow this shift dynamically, we tracked individual arrays in growing cells (Fig. 4C). Stationary-phase cells were placed sparsely between a coverslip and an agarose-hydrogel pad with fresh growth medium (see *Materials and Methods*). As cells shifted from stationary to exponential growth phase, we followed growing arrays and observed a progressive reduction in Tar occupancy in individual arrays (Fig. 4C). This shift in individual array occupancy is consistent with the overall population-level shift shown in Fig. 4B (marked by red bars). Thus, the shift in receptor occupancy between stationary and exponential growth phases observed in Fig. 4B reflects not only the formation of new arrays with altered receptor occupation but also dynamic changes of receptor occupancy within preexisting arrays. Notably, to the extent that the array structure is generally stable, a temporal shift in Tar occupancy suggests that a highly non-uniform Tar distribution can develop within the array over time.

### Sensory properties are consistent with mosaic receptor arrays

To analyze the response behavior observed in Fig. 2C, we first consider the commonly used MWC model^23, 26, 28^, applied to receptor clusters composed of both Tar (*n*_*Tar*_) and Tsr (*n*_*Tsr*_) receptors^39^ (see *Materials and Methods*). Characterizing the receptor composition in the cluster by the total number of receptors *n*_*tot*_ and the relative Tar content *r*≡ *n*_*Tar*_/*n*_*tot*_, we note that under saturating concentrations of either aspartate or serine, the inherent cooperativity in the cluster^16, 25^ leads to a sharp dependence of kinase activity on *r* (Figs. 5A and S8, and *Materials and Methods*: Eqs. 2–3). The asymmetry between aspartate and serine shown in Fig. 5A results from the inherently distinct signaling properties of Tar and Tsr (see refs. 18, 24, 25 and Fig. S9).

**Figure 5.**
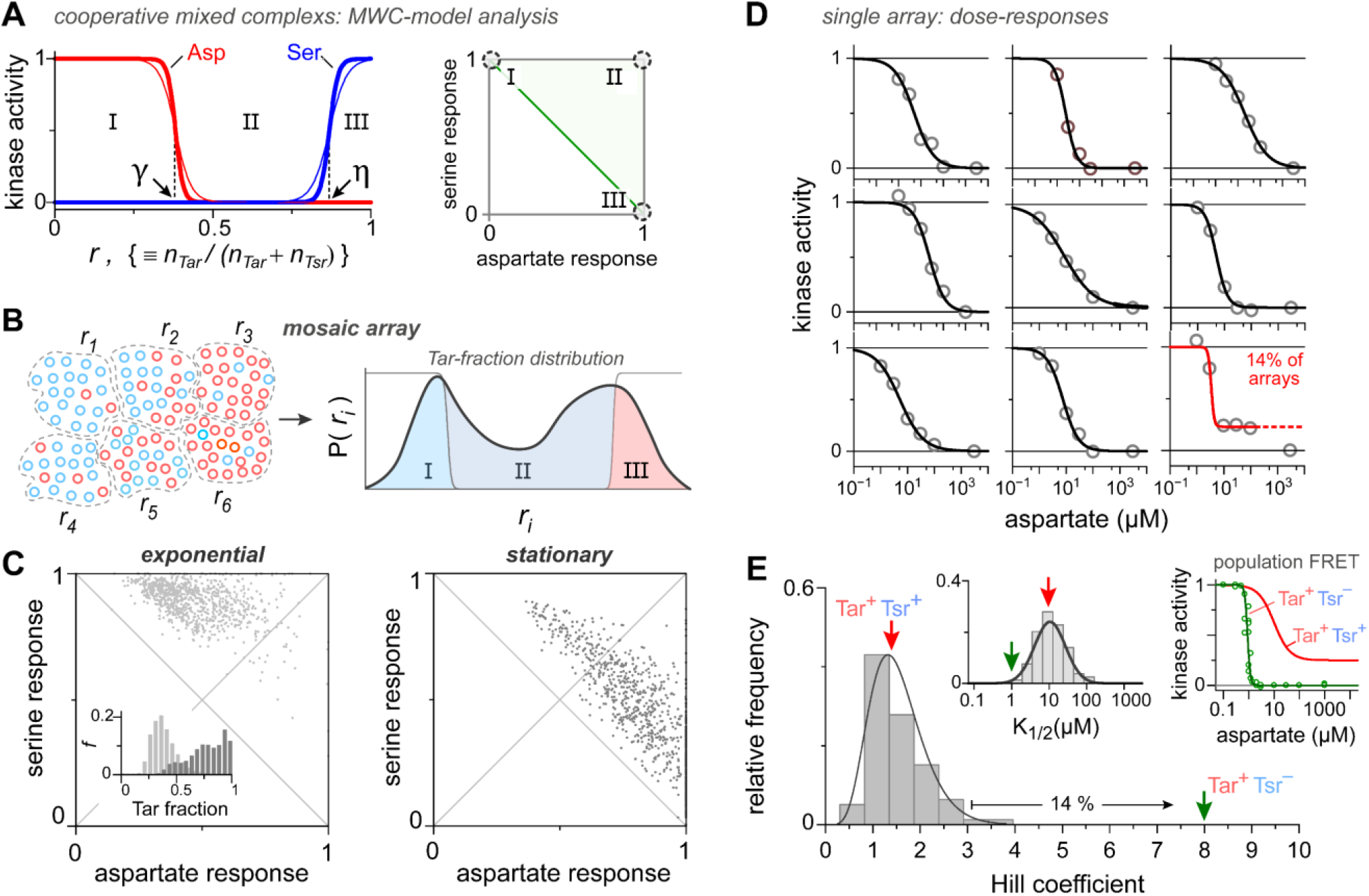
Mosaic arrays. (**A**) MWC-model-based analysis of the correlation between serine and aspartate responses (Eqs. 2–3 in Materials and Methods). A receptor cluster containing both Tar (n_Tar_) and Tsr (n_Tsr_) is considered in the presence of saturating aspartate (red lines) or serine (blue lines). The kinase ‘on’ probability is plotted as a function of the Tar fraction, *r = n*_*Tar*_ */ n*_*Tot*_, using the following parameters: *n*_*Tot*_ *= 12* (thick lines) or 6 (thin lines)^16, 25^; *K*_*on*_ */ K*_*off*_ = 150 for Tar^24,25^ and 2,700 for Tsr (Fig. S9); and *ΔE* = –0.8 for Tar and –2.6 for Tsr (in units of k_B_T). Two critical points can be defined: *r = γ* and *r = η*, where *P*_*on*_*(asp)* or *P*_*on*_*(ser)*, respectively, equals 0.5. Although exact values of these points depend on the parameters, the fundamental division only relies on the cooperative nature of the responses (see Fig. S8). Mapping these regions onto the Ser–Asp plot (right illustration) suggests that most clusters are expected to reside near the corners of the plot (dashed circles). (**B**) Schematic of a mosaic array composed of non-uniform cooperative sub-regions (dashed line), each with a distinct receptor composition, characterized by *r*_*i*_. The sensory responses of such an array are determined by the distribution of *r*_*i*_ values, and in particular by how these values are partitioned among the three regions (right schematic). (**C**) Simulated arrays (see Materials and Methods) exhibit response patterns similar to those measured experimentally (Fig. 2C), with only a moderate shift in overall Tar fraction, shown in the inset (*r*_*avg*_ ∼ 0.4 and 0.76, for the exponential and stationary phase, respectively). See also Fig. S10. (**D-E**) Single array dose-responses exhibit low cooperativity. *Part D*: Examples of dose-dependent responses, measured from individual arrays in stationary-phase VF17 cells (OD 3.5), with black lines representing fits to a Hill function. Responses are relative to the saturation value for each array. *Part E*: Distributions of the Hill coefficient and K_1/2_ characterizing the single-array responses (gray bars; 105 arrays). Approximately 14% of the arrays exhibit behavior consistent with a partial response with Hill coefficients potentially larger than 3; an example is shown in the lower-right plot in part D. The Hill coefficient and K_1/2_ values obtained from fitting the population FRET dose-responses are also indicated by arrows, and the corresponding dose-responses are shown in the inset: *red line and arrows* – Tar^+^Tsr^+^ (VF17) cells (from Fig. 2D); *green line and arrows* – Tar^+^ Tsr^-^ (MK32) cells.

The critical *r*-values at which the kinase activity is inhibited by aspartate (*γ*) or serine (*η*) define three regions (Fig. 5A): (I) clusters with *r* < *γ*, where the kinase output can only be inhibited by serine; (II) clusters with *γ* < *r* < *η*, where the kinase output can be inhibited by either serine or aspartate; and (III) clusters with *r* > *η*, where the kinase output can only be inhibited by aspartate. This analysis shows that a single MWC-like cooperative cluster typically belong to one of these regions, and thus maps to one of the three corners of the Ser–Asp response plot (Fig. 5A, right), in contrast to the broad-range responses observed in Fig. 2C.

On the other hand, diverse single-array responses can still be compatible with substantial response cooperativity if the arrays have non-uniform composition, as suggested by Fig. 4B-C. We therefore next considered mosaic-like arrays comprising multiple subregions, each with distinct receptor compositions (Fig. 5B). Assuming an array with *N* sub-regions, each with distinct Tar fraction *r*_*i*_ and maximal kinase activity *a*_*i*_, the total kinase activity is given by:

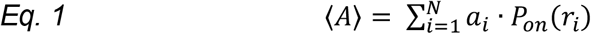

where *P*_*on*_(*r*_*i*_)is the ligand-dependent probability of subregion *i* to be in the active state. Assuming that each sub-region is still highly cooperative and well-described by the MWC model, the response properties of such mosaic arrays would essentially depend on the distribution of *r*-values within the array (Fig. 5B; see also Materials and Methods). In this case, the aspartate response approximately corresponds to the fraction of subarrays with *r*-values in regions II and III, and the serine response to the fraction of subarrays with *r*-values in regions I and II.

Thus, provided that the *r*-value distribution in the array can be sufficiently broad to encompass all three regions, arrays can occupy any location on or above the ‘additivity’ line in the Ser–Asp space. To test this proposition, we simulated array populations assembled under varying Tar and Tsr expression conditions (time-dependent *r*_*i*_ distributions) corresponding to the ‘exponential’ and ‘stationary’ phases (see Materials and Methods). Indeed, we find that even a moderate change in the average Tar fraction (as observed in Fig. 4B) can produce a significant shift in the response properties comparable to those observed in Fig. 2C (Figs. 5C and S10).

### Single-array dose-response behavior

The population-level dose-responses of VF17 cell with mixed receptor arrays (Fig. 2D), generally exhibited lower cooperativity than that observed in arrays composed of a single receptor type^16, 23, 25^. Such low cooperativity could, in principle, result from K_1/2_ diversity among individual arrays arising from differences in their composition. In this case, arrays could each retain high intrinsic cooperativity yet differ in sensitivity (K_1/2_). Alternatively, under the mosaic view of arrays (Fig. 5B), the internal diversity in receptor composition, and thus in K_1/2_, would be expected to produce low-cooperativity dose-responses even in individual arrays.

To test these propositions directly, we measured dose-dependent responses of single arrays (Fig. 5D–E), focusing on the aspartate response in stationary-phase cells, which showed especially low cooperativity (Hill coefficient ∼ 1.4) and larger responses than those in the exponential phase (Fig. 2D). Single-array responses were measured in VF17 cells as before (Figs. 1–2), except that cells were stimulated with increasing concentrations of aspartate (see *Supplementary Methods*). Several examples are shown in Fig. 5D, with each plot corresponding to a different array. The responses of each array were fitted with a Hill function (black lines), and the distributions of the resulting Hill coefficients and K_1/2_ values are shown in Fig. 5E (gray bars). Additionally, Fig. 5E (right inset) shows the population-level (FRET) dose-response behavior of the same VF17 cells (red line; Tar+Tsr+, taken from Fig. 2D) and of similar cells but lacking the Tsr receptor (green symbols; Tar+ Tsr-, MK32). The latter are expected to have more uniform array composition and indeed exhibit substantially higher cooperativity. The corresponding Hill-fit parameters for these population-level responses are indicated in the main plot as red and green arrows.

The cooperativity of most single arrays (Fig. 5E, gray bars) is comparable to that observed at the population level (red arrow) and significantly smaller than the Hill coefficient characterizing cells with more uniform arrays (Tsr^-^ cells; green arrow). The dose-response behavior of approximately 14% of the arrays is potentially consistent with higher cooperativity, but this applied mostly to only partial kinase inhibition (see Fig. 5D, lower-right plot); however, we cannot evaluate their actual cooperativity, which would require a much denser concentration scan. The overall low cooperativity observed here at the single-array level is generally consistent with the proposed mosaic organization of receptor arrays.

### Diverse sensory responses can provide an advantage in complex environments

Previous studies using distinct serine and aspartate sources have shown an increased bias toward aspartate at later stages of growth^35^. This bias was primarily dependent on the sensitivity of the cells to each effector at very low concentrations. To further test possible behavioral consequences of the diversity and bias shift observed in the responses of the arrays (Fig. 2C), we used wild-type RP437 cells, and examined how their growth phase influences their ability to overcome a repellent barrier. The barrier was created and controlled by a single source containing both MeAsp (attractant, sensed by Tar) and indole (repellent, sensed by Tsr) at the end of a long channel filled with bacterial cells (Fig. 6A)^40^. The combined effect of these effectors produces a region of effective repulsion that propagates away from the source, followed by an attractant region lagging behind. The effective strength of the repellent barrier could be modulated by the attractant (MeAsp) concentration in the source (Fig. 6B)^40^. As the repulsion lobe propagates through the channel, cells that respond effectively to the repellent (indole, Tsr) are pushed away from the source, whereas cells that respond weakly to the repellent or more strongly to the attractant (MeAsp, Tar) are more likely to cross the barrier and reach the attractant region. Indeed, previous studies have shown that Tsr^−^ cells exhibited only attractant responses, whereas Tar^−^ cells exhibited a stronger repellent response^40^.

**Figure 6.**
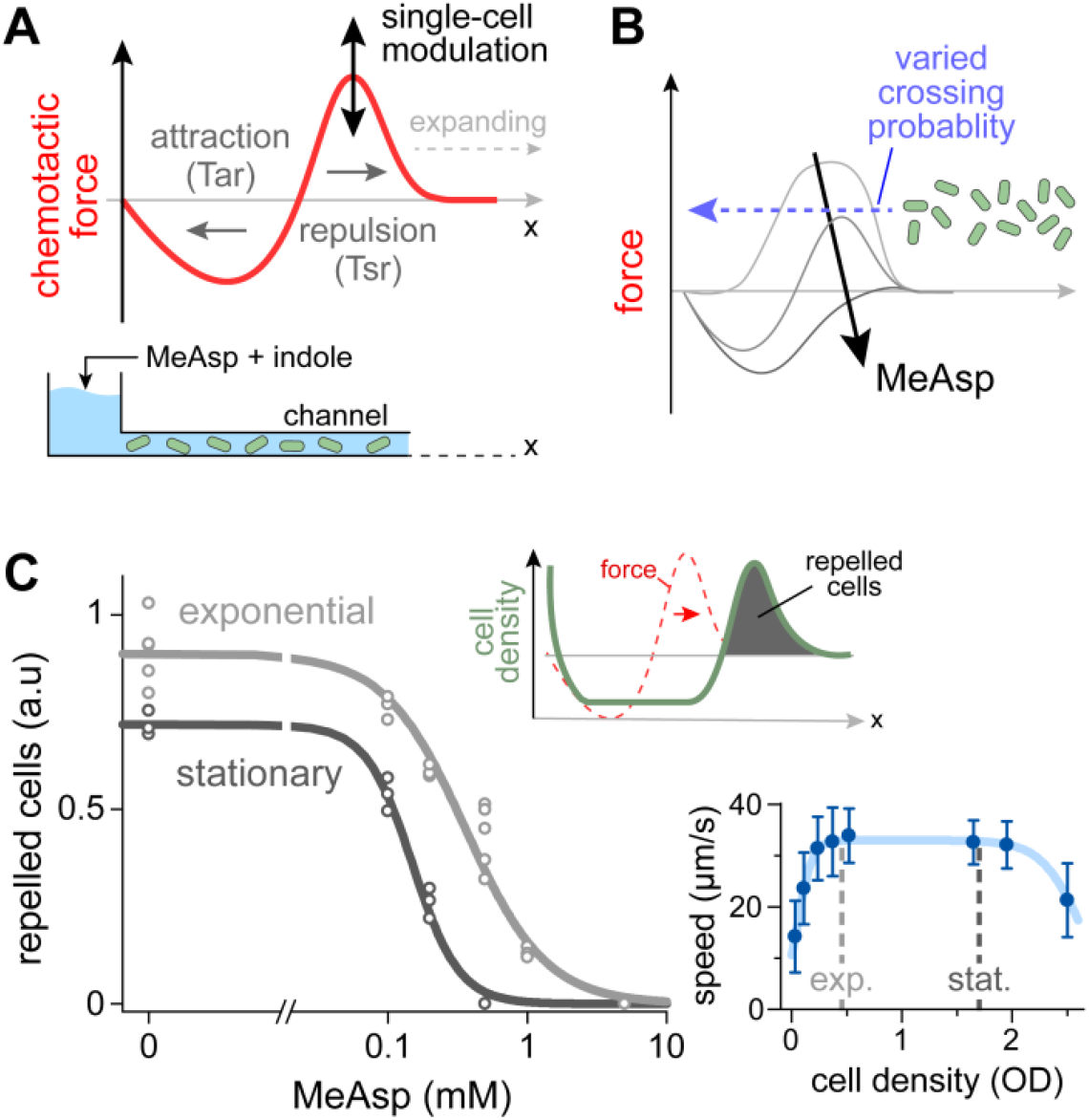
Growth-phase-dependent chemotaxis behavior in a complex effector field. (**A**) Schematic of the experimental setup. A uniform cell suspension was injected into a long (44 mm) channel. A source containing both indole (Tsr-mediated repellent) and MeAsp (Tar-mediated attractant) was then introduced at one end of the channel and allowed to diffuse into the channel (see Materials and Methods). This source creates a repellent-dominated region that expands into the channel, followed by a lagging attractant-dominated region (see ref. 40). Due to sensory variability between cells, each cell is expected to experience a modulated chemotactic force. (**B**) Addition of MeAsp modulates the magnitude of the repellent force, thereby affecting the probability of crossing toward the source. (**C**) The number of cells within the repellent lobe (inset), evaluated by the integrated fluorescence intensity three hours after introducing the source (main plot). Results are shown for GFP-expressing wild-type (RP437) cells, grown to exponential phase (OD 0.4; light gray) or early stationary phase (OD 1.7; dark gray). Each point represents an independent experiment; lines are a guide to the eye. The corresponding swimming speeds of cells at different growth stages are shown in the lower-right inset (blue symbols); error bars represent standard deviation (more than 200 cells per point).

We now examined how the bacterial growth phase influences their ability to overcome the repellent barrier for different concentrations of (MeAsp) attractant at the source. The number of cells repelled from the source at different MeAsp concentrations is shown in Fig. 6C. The swimming speed and tumble-bias of these cells at the two growth stages were generally similar (Figs. 6C and S11). We found that exponential-phase cells, with mostly Tsr-dominant array responses (Fig. 2C), were less able to overcome the repellent lobe and therefore accumulated in greater numbers away from the source (Fig. 6C). Similarly, early-stationary-phase cells, in which most arrays exhibited substantial Tar-mediated responses (Fig. 2C), were more sensitive to MeAsp addition and exhibited a sharper response to MeAsp. Overall, although cell behavior is clearly influenced by additional factors, most notably adaptation, the observed behaviors are generally consistent with the sensory properties shown in Fig. 2C.

## Discussion

By combining the kinase and the response-regulator proteins of the chemotaxis system into a hybrid protein embedded in the sensory array, we were able to measure the kinase activity of individual arrays (Fig. 1). Applied here to study arrays with native receptor compositions in cells lacking the adaptation enzymes, this method revealed highly diverse and growth-phase-dependent sensory properties (Fig. 2), with the response repertoire shifting from serine-biased in the exponential phase to aspartate-biased in the stationary phase. In both cases, however, the orthogonal stimulus was still able to inhibit array kinase activity to a widely varying degree (Fig. 2C). Analysis of these sensory responses (Fig. 5A–C), together with the low cooperativity observed in single-array dose-response measurements (Fig. 5D–E) and the dynamic receptor occupancy in individual arrays (Fig. 4B–C), suggest that arrays exhibit a non-uniform receptor organization and function as a mosaic of cooperative regions. Furthermore, the weak correlations between different arrays within the same cell (Fig. 3), together with the shift in Tar and Tsr occupancy distribution (Fig. 4B) and, in particular, the temporal variations in receptor occupancy during growth (Fig. 4C), suggest that the sensory properties of individual arrays reflect their unique assembly history (Fig. 7). This history may reflect both global temporal variations in Tar and Tsr occupancy^35, 37^ (Fig. 4) and cell-to-cell variations within the specific lineage of cells in which each array was assembled.

**Figure 7.**
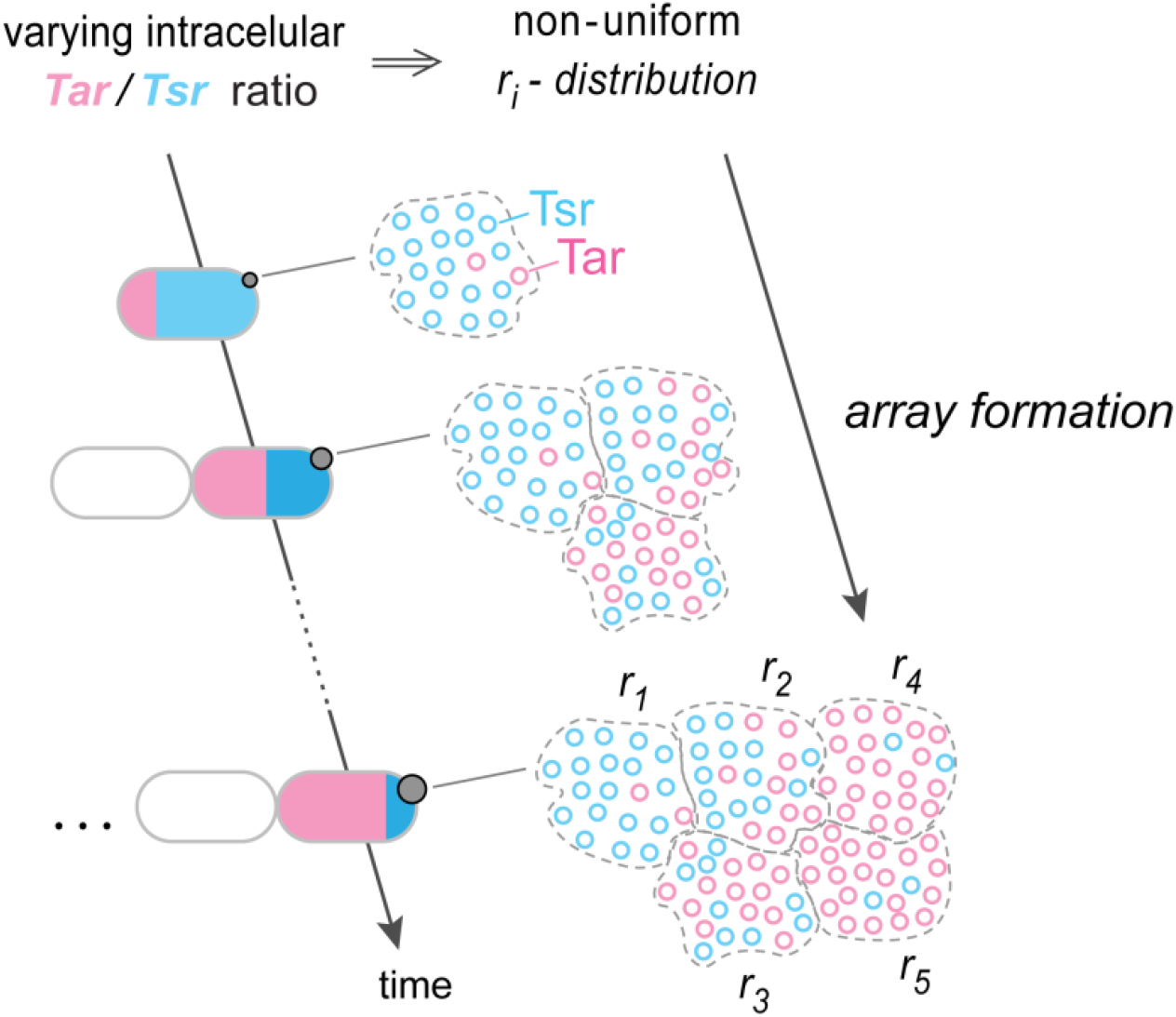
Formation of non-uniform, mosaic arrays. Two properties of receptor arrays can lead to diverse local receptor compositions (i.e., a wide distribution of *r* values): (i) Array growth is slow and spans multiple cell cycles (see Fig. 4 and ref. 11), resulting in array growth under temporal and cell-to-cell variation in Tar and Tsr expression levels (see Fig. 4A–B). (ii) Receptor arrays remain structurally stable over time, preserving earlier receptor incorporation patterns (see Discussion).

In general, mosaic multicomponent assemblies can be expected when (i) they are stable or undergo slow turnover with limited internal molecular mixing, and (ii) their components exhibit dynamic changes in expression over the timescale of their assembly. Under these conditions, the internal composition of such assemblies is shaped by their assembly history and by the expression dynamics of their components within the specific cell lineage in which they form (Fig. 7). In chemoreceptor arrays, receptor expression indeed changes dynamically over time (Fig. 4B–C)^38^ and their assembly process occurs over several generations (Fig. 4C)^11^. Several observations support the overall stability of these arrays. Evidently, arrays remain intact even under stimulation^29, 41^(Fig.1D, CheA::CheY–mYFP fluorescence). Moreover, in line with their highly structured hexagonal molecular organization (Fig. 1), crosslinking experiments have demonstrated very limited mixing of receptors within arrays^42^. Homo-FRET measurements have revealed slow conformational dynamics in receptor arrays following exposure to attractant effectors^43^; however, these changes likely reflect local receptor conformational changes rather than global reorganization. Additionally, using tagged receptors, limited and slow fluorescence recovery was measured after photobleaching^44^, but it remains unclear to what extent this recovery indeed reflects internal mixing within the array. Thus, even if some internal dynamics occur, they are unlikely to mix the receptor population sufficiently to achieve uniformity.

Population-level measurements of cells expressing both Tar and Tsr typically exhibit lower cooperativity than those of uniform arrays in cells expressing either receptor type alone^23^. Low cooperativity responses were also evident here in population-level measurements of cells with native receptor compositions (Fig. 2D). Low Hill-coefficient values were also observed in previous measurements of single-cell responses^31^. As shown in Fig. 5D–E, this reduced cooperativity persists even at the single-array level. Notably, the apparent cooperativity of these arrays is comparable to that observed with dispersed (non-clustered) core complexes, even though networking of core complexes substantially increases cooperativity^16, 18, 19^. Such low cooperativity in mixed arrays is generally consistent with the mosaic view proposed in Fig. 5B, in which arrays retain their inherent molecular properties, including local conformational coupling within or between core complexes, while their non-uniform composition introduces internal K_1/2_ variability that ultimately reduces the apparent cooperativity. Moreover, given the intricate conformational coupling within core complexes^17, 18^, the way in which different receptor types are distributed even within core complexes could also influence the overall response.

Interestingly, in mosaic arrays, signal integration is expected to depend on the size of the cooperative sub-regions; larger cooperative regions would average out more of the underlying non-uniformity, thereby modulating their *r*-values. Such changes would strongly influence the correlation between the array responses to different signals. For example, an array containing two sub-regions with *r*-values *0*.*1* (mostly Tsr) and *0*.*9* (mostly Tar) would display additive behavior (Fig. 5B), whereas combining these two sub-regions into a single region with an *r*-value of *0*.*5* would produce an array that can be fully inhibited by either stimulus. Indeed, dynamic changes in the size of the cooperative regions have been previously suggested^45^. Additionally, temporal switching of kinase-activity was observed in single cells with uniform arrays tuned to a low-activity state^31, 46^, suggesting that, at least under certain conditions, cooperative regions may substantially expand. Moreover, adaptation, by lowering the activity state of the receptors in the absence of ligand, may shift the critical points in Fig. 5A further apart, thereby expanding the middle zone (Fig. 5A and S8) and promoting full responses to multiple stimuli. Adaptation may also reduce the diversity in ligand sensitivity (K_1/2_) among different regions of the array^47, 48^, potentially enhancing their overall apparent cooperativity.

Finally, mosaic arrays allow substantial diversity in the fraction of kinase activity controlled by each sensor type within the array and, consequently, in the overall array responses, leading to diversity in cellular behaviors. Such diversity was indeed observed in single cell responses^31^, and can be advantageous at the population level. For example, it can facilitate coordinated chemotaxis behaviors^32, 49^, or, as shown in Fig. 6, allow subset of cells to overcome chemotactic barriers created in complex environments, thereby promoting bet-hedging type of behavior. Specifically, cells that strongly respond to the Tsr-mediated repellent would avoid the repellent environment but also fail to reach the attractant source, whereas, cells that respond weakly to the repellent or more strongly to the Tar-mediated attractant would be able to cross the repellent region and reach the source.

## Materials and Methods

### Bacterial strains and plasmids

Strains used here are derivatives of *E. coli* K-12 strain RP437(eda+)^50^. Their relevant genotypes are as follows: VF17 [*cheA(ΔP2)::cheY*–*myfp*, Δ(*cheR, cheB, cheY), cheZ(F98S)*–*mscarle-I*]; MK27 [*cheA(ΔP2)::cheY*–*myfp*, Δ*(cheR, cheB, cheY, cheZ), tsr*–*mscarlet-I*]; MK30 [*cheA(ΔP2)::cheY-myfp*, Δ(*cheR, cheB, cheY, cheZ), tar-mscarlet-I*]; and MK32 [*VF17+ tsr::ccdB-kan* ]. We also used strain UU2828 *[cheR, cheB, cheY, cheZ]* expressing Che-mCherry *and* CheZ(F98S)-mYFP from pKG110 vector (pAV109; cam^R^, NaSal 0.8 μM). We used the mScarlet*-I* variant throughout this work.

### Strain construction

Strain VF15 was created from strain *UU2828* by replacing the *cheA*.*P2* (158–230) region with *cheY-myfp* and introducing the *M98L* point mutation in *cheA* to prevent expression its short variant^34^. Strain VF17 was then created by inserting back *cheZ(F98S)-mscarlet-I*, along with its native RBS (20 bp), 18 bp downstream of *tap*. Strains MK27 and MK30 were constructed from strain VF15 by replacing respectively *tsr* with *tsr-mscarlet-I* or *tar* with *tar-mScarlet-I*. Strain MK32 was created by replacing tsr 211-551 with ccdB-Kan cassette.

### Media and growth conditions

Cells were pre-grown overnight to saturation in TB medium (10 g/l Bacto-Tryptone and 5 g/l NaCl) at 30°C. Cells were then diluted in 10 mL of M9 minimal medium with 1% sodium gluconate and supplemented with 1 mM MgCl_2_, 1 mM Na_2_SO_4_, 30 μM vitamin B_1_, and a mixture of the amino acids (Histidine, Threonine, Methionine, and Leucine, 1 mM each), and allowed to grow overnight with shaking at 32°C to reach stationary cultures at optical density (OD) of 3.5–4. Overnight cultures were then diluted to an OD of 0.035 in fresh minimal media and grown to the desired OD at 32°C. Cells were harvested at the indicated OD and washed twice with motility buffer (10 mM potassium phosphate, 0.1 mM EDTA, 1 μM methionine, 10 mM lactic acid, pH 7.1) before imaging. For growth experiments (Fig. 4C), cells were taken directly from the overnight culture and allowed to grow on an agarose hydrogel (1.2%) made from fresh minimal media. In experiments involving UU2828/pAV109 cells (Figs. 1C and S2), following the initial overnight growth in TB, cells were diluted 1:100 in TB and allowed to grow to OD 0.45.

### Imaging

Fluorescence images were obtained using a Nikon-Ti inverted microscope equipped with a 100× Plan-Fluor objective (1.3 NA), a xenon lamp (Sutter Instruments), and a camera (Andor Technology), and filter sets: Ex-ET560/40–EmET630/75 (red channel) and ExET490/20– EmET353/50 (yellow channel). Cells were cultured as described above. To measure kinase activity response, cells were placed between a 1.2% agarose hydrogel pad and a poly-L-lysine-coated coverslip for 20 minutes before transferring the coverslip to a flow chamber. Initially, cells were in motility buffer, and an image of the unstimulated state was taken. They were then exposed to different stimuli for 1 minute before imaging and washed with buffer between exposures. A final image was taken after returning the cells to buffer. In the single array dose response experiments (Fig. 5D-E), cells were exposed to increasing concentrations of aspartate for 1 minute before imaging. See *Supplementary Methods* for detailed description of the analysis. In order to evaluate receptor occupation (Fig. 4), cells were placed between a 1.2% agarose hydrogel pad and a coverslip.

### Population FRET measurements

The kinase activity, averaged over cell populations, was measured by following FRET between tagged CheY and CheZ proteins^20, 22^. Flow cell assembly and FRET measurements were done as described before^19, 43^. Activity-dependent FRET occurs between CheA::CheY–mYFP and CheZ–mScarlet-I in the newly constructed strains (VF17 and MK32), and between cheY-mCherry and CheZ–mYFP expressed from plasmid pAV109 (pKG110) in strain UU2828.

### Image analysis

Images were analyzed using custom-written software to quantify changes in CheZ localization to the arrays, a proxy for kinase activity responses, and were manually confirmed in several cases. For each array, the maximum intensity (*I*_*array,i*_) and the mean intensity in the cell body outside array regions (*I*_*body,i*_) were measured relative to intensity outside the cell in both the ‘red’ (CheZ-mScarlet) and yellow (CheA::CheY-mYFP) channels, with *i* indicating different stimulation conditions. The ‘yellow’ channel provided information about the size and stability of the arrays under the various conditions. To obtain the kinase responses, intensities from the red channel were first corrected for photobleaching according to:

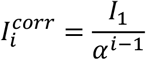

where *α* ranged from 0.80 to 0.87 (see Fig. S3) and *i* denotes the image sequence number. The added noise from this correction was estimated to be approximately 2–5% (Supplementary Text 2). The peak amplitude of each array (Fig. 1D) was calculated as the difference between the measured array peak intensity and the background intensity at the array position contributed only by the unbound CheZ:

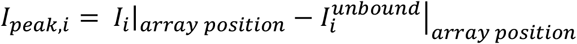

where the background intensity was determined as

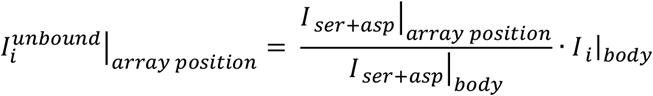

Changes in peak intensity upon stimulation with either aspartate or serine were then defined as:

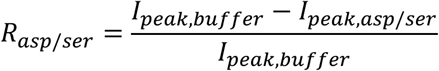

Additional source of noise (∼10%) arises primarily when the response approaches 1, namely, when no clear peak can be identified under stimulation conditions (i.e., *I*_*peak,i*_∼0). In these cases, the measured intensity at the array position, *I*_*i*_, closely matches the background intensity at this position due to unbound CheZ, 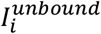, making their difference sensitive to uncertainties in identifying of the exact peak position. This effect accounts for the observed deviations of R values beyond 1 in Fig. 2C. For more details see Supplementary Methods.

### MWC model for mixed clusters: dependence on Tar fraction

Following ref. 39, we considered highly cooperative receptor clusters composed of *n*_*tot*_ receptors: Tar (*n*_*Tar*_) and Tsr (*n*_*Tsr*_). The kinase ‘on’ probability is then given by:

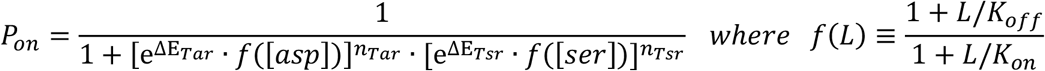

Or, by defining *r*≡ *n*_*Tar*_/*n*_*tot*_ :

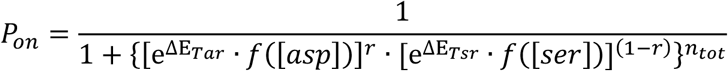

Assuming that for *L* >*K*_*on*_, we approximate *f*(*L*)∼*K*_*on*_/*K*_*off*_, and thus:

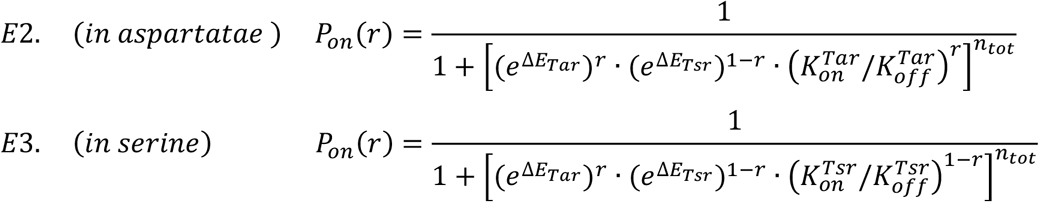

### Simulated arrays

We numerically generated arrays with non-uniform internal compositions of Tar and Tsr using distinct statistical distributions corresponding to ‘exponential’ and ‘stationary’ growth phases (Fig. S10). Receptor statistics were chosen to produce responses similar to those observed experimentally, while keeping the overall shift in the averaged Tar fraction quite moderate (Fig.5C and S10). Each growth phase was divided to three time periods, each characterized by a unique distribution of Tar-fraction *r*-values (Fig. S10). The mean *r*-value gradually shifted over time. For simplicity, normal distributions were used. During each period, new arrays were randomly nucleated, increasing the total number of arrays 4-fold in the stationary and 8-fold in exponential phases. At each time step, core units were added to both new and preexisting arrays, according to their nucleation point. For each addition, an *r*-value was randomly drawn from the corresponding distribution, and core units were added such that, on average, the ratio of Tar and Tsr reflect this value. On average, 30 core complexes were added for each choice of *r*-value, representing time correlation. At the end of each growth phase (stationary or exponential’), core unites were grouped into strongly cooperative regions, and the signaling properties of these arrays were evaluates using Eq. 1. This cycle of exponential and stationary phases, was iteratively repeated several times.

### Behavioral chemotaxis assay

The assay was performed as previously described in ref. 40 (see schematic in Fig. 6A). RP437 cells (UU2547, courtesy of Sandy Parkinson, University of Utah) expressing GFP from a plasmid (pSA11, pTrc99a, induced by 75 μM IPTG during the final growth step) were harvested at the desired growth stage, suspended in motility medium (with added NaCl 5 gr/l) to final density OD_600_ 0.1, and injected into a long channel (44 mm long, 5 mm wide, 0.2 mm high; ibidi μ-Slide), forming an initially uniform distribution. Then, one end of the channel was sealed with 1.5% (w/v) agarose hydrogel in motility buffer, and the other end (the ‘source’) was filed with 1.5% (w/v) agarose hydrogel in motility media containing 1 mM indole and varying concentrations of the non-metabolizable analogue of aspartate, α-methyl-aspartate (MeAsp). The channel was then incubated at 30°C and after 3 hours, cell redistribution was monitored by fluorescence microscopy using a 20X objective (0.5 NA). The amount of ‘repelled bacteria’ was quantified by integrating the density of cells migrating away from the source relative to the initial baseline cell density (see Fig. 6C, upper inset).

To quantify the swimming speeds at different growth stages (Fig. 6C, lower inset), cells were suspended in the same motility medium at low cell density (OD_600_ ∼ 0.03) and injected into an ibidi μ-Slide channel for observation. To induce smooth swimming and enable quantification of swimming velocity, both serine (2 mM) and MeAsp (10 mM), were added to the suspension. Time-lapse imaging was then performed at 10–15 frames per second using a 20× (NA 0.5) objective near the bottom surface and bacterial tracks were detected using the TrackMate plugin for ImageJ.

To evaluate tumbling bias (Fig. S11), cells were diluted in motility medium (OD 2·10^-4^) and placed in a μ-slide (ibidi 80326, coated with BSA). Time lapse images were captured near the top surface using a 10x (NA 0.45) objective at ∼15 frames per second for 100 seconds and analyzed in 2D^51^, using the algorithm described in ref. 52 to identify tumble events. Only tracks longer than 3 seconds were analyzed.

## Abbreviations

IPTG: isopropyl-β-D-thiogalactopyranoside
FRET: Förster resonance energy transfer

## Acknowledgments

We thank John S. Parkinson (Utah University) for careful reading of the MS and helpful comments. This work was supported by the Israeli Foundation of Sciences and Humanities, and by research grant BSF 2017356 from the U.S.-Israel Binational Science Foundation. We also thank the Milner foundation (NL).

## Notes

### Competing Interest Statement

The authors have declared no competing interest.

