## Supplementary Information for "Single-array measurements reveal non-uniform, mosaic-like chemosensory arrays in bacteria"

**Supplementary Text 1.** Estimating the interference between the kinase activity in different arrays.

**Supplementary Text 2.** Analyzing photobleaching and related the uncertainty.

**Supplementary Methods.** Single array responses.

**Supplementary Figures:** 1 – 11

**Text 1. Estimating the interference between the kinase activity in different arrays.**

The relevant phospho-transfer reactions are:

Auto-kinase:

$$\mathrm{CheA.P1} + ATP + CheA.3-5 \leftrightarrow [\mathrm{CheA.P1} \cdot ATP \cdot \mathrm{CheA.P3}-5] \rightarrow \mathrm{CheA.P1}p + ADP + CheA.3-5$$

Phosphotransfer: $\mathrm{CheA.P1}p + Y \leftrightarrow [\mathrm{CheA.P1}p \cdot Y] \rightarrow \mathrm{CheA.P1} + Yp$

Dephosphorylation: $Yp + Z \leftrightarrow [Yp \cdot Z] \rightarrow Y + Z$

Where the subscript 'p' denotes the phosphorylated form of the protein/domain. Since the global kinase activity in the cell must be balanced by the total CheZ activity, changes in the kinase activity in one receptor array can influence the inherent kinase activity of the others.

To estimate this potential interference we consider two arrays, and ask how much the kinase activity in array 1 changes upon turning off array 2.

**Before turning off array 2:** In steady state, the 'kinase activity', labeled: $A$ — the phosphorylation rate of CheA.P1 — is generically balanced by the dephosphorylation rate of CheY, $V_Z$. Moreover, since both CheA and CheY reside in the same hybrid protein within the array, this balance must occur within each array. Thus

$$E1. \qquad A_1 = V_Z(array\ 1) = [Yp \cdot Z]_1 \cdot k_{cat}^z \qquad and \qquad A_2 = V_Z(array\ 2) = [Yp \cdot Z]_2 \cdot k_{cat}^z$$

But also holds globally in the cell, where according to Michaelis-Menten we generally have:

$$E2. \qquad A_{tot} = A_1 + A_2 = (\ [Yp \cdot Z]_1 + [Yp \cdot Z]_2\ ) \cdot k_{cat}^z = [Z_{tot}] \cdot \frac{\{[Yp1] + [Yp2]\}}{K_M + \{[Yp1] + [Yp2]\}} k_{cat}^z$$

Defining $\gamma_0$ to be the fraction of CheZ bound to the arrays in a cell in the absence of any ligand, and using *E2* we find:

$$E3. \qquad \gamma_0 \equiv \frac{[Yp \cdot Z]_1 + [Yp \cdot Z]_2}{[Z_{tot}]} = \frac{\{[Yp1] + [Yp2]\}}{K_M + \{[Yp1] + [Yp2]\}}$$

The value of $\gamma_0$ can be directly estimated from the data by evaluating the change in the unbound CheZ (background 'red' fluorescence in the cell) before and after inhibiting the kinase activity by addition of both serine and aspartate, which release all the CheZ from the arrays. As seen in Fig. S1A(inset) such an estimate yield: $\gamma_0 \lesssim 0.4$.

We also define $\beta$ as:

$$E4. \qquad \beta \equiv \frac{1}{1+\{[Yp1]+[Yp2]\}/K_M} = 1-\gamma_0$$

We can express the total kinase activity (from *E2*) as:

$$E5. \qquad A_{tot} = A_1 + A_2 = \{[Z_{tot}]\cdot k_{cat}^{z}\}\cdot\gamma_0 = \{[Z_{tot}]\cdot k_{cat}^{z}\}\cdot\beta\cdot([Yp1]/K_M + [Yp2]/K_M)$$

Assuming that the total amount of bound CheZ partitions between the arrays according to their CheYp levels only, we have (from *E5*):

$$E6. \qquad A_1 = \{[Z_{tot}]\cdot k_{cat}^{z}\}\cdot\beta\cdot[Yp1]/K_M \quad and \quad A_2 = \{[Z_{tot}]\cdot k_{cat}^{z}\}\cdot\beta\cdot[Yp2]/K_M$$

Finally, defining: $\chi_{Yp1} \equiv \frac{[Yp1]}{K_M}$; $\chi_{Yp2} \equiv \frac{[Yp2]}{K_M}$; $\gamma_1 \equiv \frac{[Yp\cdot Z]_1}{[Z_{tot}]}$; $\gamma_2 \equiv \frac{[Yp\cdot Z]_2}{[Z_{tot}]}$; we can rewrite *E6* as:

$$E7. \qquad \gamma_1 = \beta\cdot\chi_{Yp1} \quad and \quad A_1 = \{[Z_{tot}]\cdot k_{cat}^{z}\}\cdot\gamma_1 = \{[Z_{tot}]\cdot k_{cat}^{z}\}\cdot\beta\cdot\chi_{Yp1}$$

$$\gamma_2 = \beta\cdot\chi_{Yp2} \quad and \quad A_2 = \{[Z_{tot}]\cdot k_{cat}^{z}\}\cdot\gamma_2 = \{[Z_{tot}]\cdot k_{cat}^{z}\}\cdot\beta\cdot\chi_{Yp2}$$

Note that $\beta$ reflects the level of interference between the arrays, such that for $\{[Yp1]+[Yp2]\}/K_M << 1$ $\beta$ approaches 1 and $A_1$and $A_2$ become independent of one another (Fig. S1A). Also note that we inherently assume here that ATP consumption by kinase activity is small compared with the total energy beget in the cell, and thus, ATP-mediated interference between arrays is not significant.

**After turning off array 2:** We now turn to estimate the change in the kinase activity in array '1' ($A_1$) upon turning the kinase off in array '2', namely, setting: $A_2 \to 0$ , $Yp \to 0$ , $[Yp \cdot Z]_2 \to 0$ . Immediately after array 2 was turned off, $\beta$ has been changed, and thus causing an imbalance in array '1', which will then reach a new equilibrium by adapting the CheYp and CheA.P1p levels, and possibly also adapting the inherent kinase activity $A_1$.

— *Hereafter we only refer to array 1: before setting* $A_2 \to 0$ *(labeled 'pre') or after (labeled 'final')* —

In the <u>final</u> steady state of array 1, from *E1-3* we get:

$$\gamma_{final} \equiv \frac{[Yp \cdot Z]_f}{[Z]_{tot}} = \frac{\chi_{Yp}^{f}}{1+\chi_{Yp}^{f}} = \beta_f \cdot \chi_{Yp}^{f} \quad and \quad A_{final} = k_{cat}^{z} \cdot [Yp \cdot Z]_f = \{[Z_{tot}] \cdot k_{cat}^{z}\} \cdot \gamma_f$$

And, from *E7*:

$$E8. \qquad \frac{A_{final}}{A_{pre}} = \frac{\gamma_{final}}{\gamma_{pre}} = \frac{\beta_{final} \cdot \chi_{Yp}^{f}}{\beta_{pre} \cdot \chi_{Yp}^{pre}} = (\beta_{final}/\beta_{pre}) \cdot \frac{\chi_{Yp}^{f}}{\chi_{Yp}^{pre}} = (\beta_{final}/\beta_{pre}) \cdot \frac{[Yp]_{final}}{[Yp]_{pre}}$$

<u>On the other hand</u>, changes in CheYp in array 1 would cause also changes in the level of CheA.P1p and, in turn, can lead to changes in kinase activity.

The phosphotransfer from CheA.P1 to CheY relies on direct interactions between them and thus:

$$E9. \qquad \frac{\Delta[P1]}{[P1]} \sim -\frac{\Delta[Yp]}{[Yp]}$$

The relation between the change in CheA.P1p and that in kinase activity, *A*, is analyzed in Box 1 (Fig. S1B). We find that

$$E10. \qquad \frac{A_{final} - A_{pre}}{A_{pre}} \sim \frac{1}{2} \cdot \frac{\Delta[P1]}{[P1]}$$

From *E9* and *E10*, we find a relation between changes in $[Yp]$ and changes the kinase activity:

$$E11. \qquad \frac{A_{final} - A_{pre}}{A_{pre}} \sim \frac{1}{2} \cdot \frac{\Delta[P1]}{[P1]} \sim -\ \frac{1}{2} \cdot \frac{\Delta[Yp]}{[Yp]} = -\frac{1}{2} \cdot \frac{[Yp]_{final} - [Yp]_{pre}}{[Yp]_{pre}}$$

Therefore:

$$\frac{[Yp]_{final}}{[Yp]_{pre}} = 3 - 2 \cdot \frac{A_{final}}{A_{pre}}$$

Inserting these relations into *E8*, we finally have (see Fig. S1C):

$$E12. \qquad \frac{A_{final}}{A_{pre}} = (\beta_f/\beta_{pre}) \cdot \left[3 - 2 \cdot \frac{A_{final}}{A_{pre}}\right] \longrightarrow \frac{A_{final}}{A_{pre}} = \frac{3 \cdot (\beta_{final}/\beta_{pre})}{1 + 2 \cdot (\beta_{final}/\beta_{pre})}$$

with

$$\beta_{pre} \equiv \frac{1}{1 + \{[Yp1]_{pre} + [Yp2]_{pre}\}/K_M} \qquad and \qquad \beta_{final} \equiv \frac{1}{1 + [Yp1]_{final}/K_M}$$

Therefore, depending on the relative size of the arrays' activity we have (see Fig. S1C):

<u>For</u> $\qquad A_1 \sim A_{tot}\ \ (A_1 \gg A_2) \longrightarrow [Yp1]_{pre} \gg [Yp2]_{pre} \longrightarrow \beta_{final} \to \beta_{pre}$

and, from *E12:*

$$\frac{A_{final}}{A_{pre}} = 1$$

<u>For</u> $\qquad A_1 \ll A_{tot}\ \ (A_1 \to 0) \longrightarrow [Yp1]_{final} \ll K_M \longrightarrow \beta_{final} \to 1$

and, from *E12 and E4:*

$$\frac{A_{final}}{A_{pre}} = \frac{3}{2 + \beta_{pre}} \xrightarrow{for\ \ \gamma_0 \sim 0.4} 1.15$$

Thus, the kinase activity changes in array 1 can be expected to be typically 10% and can reach ~ 15% for very small arrays, where $A_1 \to 0$ .

**Box 1 (**Fig. S1B)

To roughly estimate the sensitivity of the kinase activity ($A$) to changes in the reactant [P1], while keeping the level of ATP constant, we start from the following Michaelis–Menten approximation:

$$A \sim V_{max} \cdot \frac{[P_1] \cdot [ATP]}{K_{p1} \cdot K_{ATP} + K_{ATP} \cdot [P_1] + K_{p1} \cdot [ATP] + [P_1] \cdot [ATP]}$$

By defining: $\chi_{p1} \equiv \frac{[P_1]}{K_{p1}}$ and $\chi_{ATP} \equiv \frac{[ATP]}{K_{ATP}}$ , we can rewrite the kinase activity as:

$$A \sim V_{max} \cdot \frac{\chi_{p1} \cdot \chi_{ATP}}{1 + \chi_{p1} + \chi_{ATP} + \chi_{p1} \cdot \chi_{ATP}}$$

Thus, the derivative of the kinase activity with respect to [P1] is therefore:

$$\frac{\partial A}{\partial [P_1]} = \frac{\partial A}{\partial \chi_{p1}} \cdot \frac{\partial \chi_{p1}}{\partial [P_1]} = V_{max} \cdot \frac{\chi_{ATP} \cdot \left(1 + \chi_{p1} + \chi_{ATP} + \chi_{p1} \cdot \chi_{ATP}\right) - \left(1 + \chi_{ATP}\right) \cdot \chi_{p1} \cdot \chi_{ATP}}{\left(1 + \chi_{p1} + \chi_{ATP} + \chi_{p1} \cdot \chi_{ATP}\right)^2} \cdot \frac{1}{K_{p1}}$$

$$= V_{max} \cdot \frac{\chi_{ATP} \cdot (\chi_{ATP} + 1)}{\left(1 + \chi_{p1} + \chi_{ATP} + \chi_{p1} \cdot \chi_{ATP}\right)^2} \cdot \frac{1}{K_{p1}}$$

Since [ATP] or [CheA.P1] are thought to be close to their respective equilibrium dissociation constants (ref. 33), we evaluate this derivative near $\chi_{p1} \sim 1$ and $\chi_{ATP} \sim 1$, yielding:

$$\frac{\partial A}{\partial [P_1]} \sim \frac{V_{max}}{8 \cdot K_{p1}} \quad \text{while} \quad A \sim \frac{V_{max}}{4}$$

Therefor

$$\frac{\Delta A}{A} \sim \frac{1}{A} \cdot \frac{\partial A}{\partial [P_1]} \, \Delta [P_1] = \frac{1}{2} \cdot \frac{\Delta [P_1]}{K_{p1}} \xrightarrow{[P_1] \sim K_{p1}} \sim \frac{1}{2} \cdot \frac{\Delta [P_1]}{[P_1]}$$

**Text 2. Analyzing photobleaching and related the uncertainty** (Fig. S3)**.**

When calculating the normalized response to serine, we compute:

$$R_{ser} = \frac{(\Delta\, red/yellow)_{ser}}{(\Delta\, red/yellow)_{ser+asp}} \quad \text{and} \quad R_{asp} = \frac{(\Delta\, red/yellow)_{asp}}{(\Delta red/yellow)_{ser+asp}}$$

where $\Delta red$ marks the intensity shift of the 'red' fluorophore (CheZ-mScarlt) peak amplitude (Fig. 1D), $yellow$ marks the peak amplitude of the 'yellow' fluorophore (CheA::CheY-mYFP), and the subscript denotes the applied stimulus. The yellow fluorophore, which reports of potential changes in the array itself, did not exhibit substantial bleaching. On the other hand, bleaching of the red fluorophore was noticeable, and approximated by the fractional change in fluorescence per image, and thus for a series of images:

$$I_{measured}(n) = I_0 \cdot \alpha^{n-1}$$

where $n$ is the image number within the sequence, see examples in Fig. S3A.

To correct for bleaching, we divided the measured intensities by $\alpha^{n-1}$:

$$I(n) = I_{measured}(n)/\, \alpha^{n-1}$$

Thus, inaccurate $\alpha$ may also lead to certain inaccuracy in the normalized responses, $R_{ser}$ and $R_{asp}$. To estimate the uncertainty in $R_{ser}$ and $R_{asp}$ ($\Delta R$) due to uncertainty in $\alpha$ ($\Delta\alpha$), we calculated:

$$\Delta R^{-}(\alpha; \Delta\alpha) = |R(\alpha) - R(\alpha - \Delta\alpha)|$$

$$\Delta R^{+}(\alpha; \Delta\alpha) = |R(\alpha) - R(\alpha + \Delta\alpha)|$$

For the purpose of bounding the uncertainty, we define

$$\Delta R(\alpha; \Delta\alpha) = \max{(\Delta R^{-}, \Delta R^{+})}$$

Notably, consistent with the small uncertainty in $\alpha$ ($\Delta\alpha/\alpha \approx 2.5\%$; Fig. S3B), the resulting $\Delta R(\alpha; \Delta\alpha)$ was smaller than 2% for both serine and aspartate (Fig. S3C).

**Supplementary Methods: single array responses.**

Each experiment consists of a series of images taken under distinct conditions. At every condition, three images were taken: phase contrast (phase), yellow fluorescence (yellow), and red fluorescence (red). See *Materials and Methods*. Prior to their analysis, images were smoothed twice using 3x3 running average. Images were then analyzed using a dedicated software running on Jupyter notebook, allowing semi-automatic analysis with close supervision.

The effectively 1D response analysis shown in Fig. 1D illustrates the principles of the approach used here and could applied in limited cases. However, the general analysis was carried out in two dimensions (2D), as follows:

The analysis can be conceptually divided into two steps:

**Step 1: Segmentation.** Initially, a small group of cells (~ 1-3 cells) was selected, and based on the phase image, a local thresholding scheme is used to distinguish the cells from their background, and their contour is detected using the OpenCV library. Arrays are then detected based on the yellow channel using a local thresholding scheme and their contour was detected. Both thresholding steps (phase and yellow) were done by manual selection of the local threshold; however, since only the peak intensities of each cluster were used, the responses and thus the threshold value did not affect the analysis.

**Step 2: Data extraction.** Each cell was separately analyzed. For each array within this cell, the sequence of images taken in the yellow channel were first analyzed and peak yellow intensity (in 2D) and its location, were recorded. The analysis of the dynamic red channel was more subtle. First, using the initial red image (in buffer) a local (2D) maximum was searched within the area defined by the initial segmentation in the yellow channel, and its value and location were recorded. The shift in the peak location between the initial red and yellow channels was also recorded. Similar procedure was used to record the red peaks in the following images, under different stimuli. However, in some of these images the red peak diminished (reflecting low kinase activity), and thus, no local maximum could be detected. In those cases, the expected array location was evaluated based on the position of the array in the yellow channel under the same conditions and the shift found between the red and yellow peaks recorded in the initial (buffer) image. The intensity (red) in this array location was then recorded.

Additionally, at each step, a 'background' intensity was recoded outside the cell (averaged over 9x9 pixels), and cell 'baseline' was recorded within the cell, along a rectangle at the center of cell (3 pixels in width).

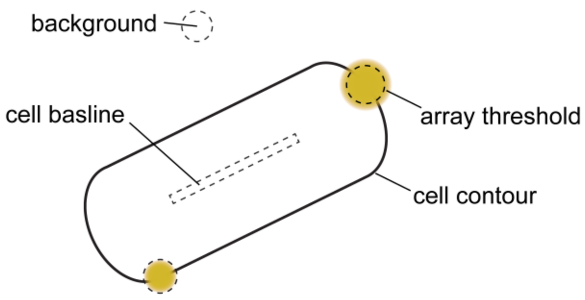


Part of arrays were excluded from this analysis based on the following general criteria:

- Cells that were in physical contact.
- Arrays that their edges overlapped or were too close to be clearly distinguished.
- Arrays that were too small to be clearly distinguished, or were not stable.

**Quantifying the responses.** Array responses were then calculated as follows.

First, the intensity outside the cell was subtracted from all recorded intensity values, that were then corrected for bleaching (see Materials and Methods, Fig. S3). Then, the peak (red) amplitude of each array during stimulus $i$ (Fig. 1D) was calculated as the difference between the measured maximum red intensity (CheZ-mScarleti) inside the array contour — $I(array)_i$ — and the intensity in the array location contributed only by the unbound CheZ — $I(array)_i^{unbound}$

$$I(peak)_i = I(array)_i - I(array)_i^{unbound}$$

To account for the fact that the amount of unbound CheZ changes upon stimulation, as part of it bind to the array, the actual expected background intensity at the array location $I(array)_i^{unbound}$ was determined, based on the local (red) intensity recorded in the presence of both stimuli (serine and aspartate) — $I(array)_{ser+asp}$ — as follows:

$$I(array)_i^{unbound} = \frac{I(array)_{ser+asp}}{I(body)_{ser+asp}} \cdot I(body)_i$$

Here, $I(body)_i$ denotes the 'cell baseline' described above.

Responses were then calculated as the fraction of kinase activity that was turned off by a specific stimulus '*i*' :

$$R_i = \frac{I(peak)_{buffer} - I(peak)_i}{I(peak)_{buffer}}$$

where $I(peak)_{buffer}$ is the average of the red peak intensity in buffer.

Additional examples for such responses are shown in Fig. S4.

**Uncertainty estimate.** In the zero-kinase state, the array intensity $I(array)_{ser+asp}$ lacks a distinct peak and was therefore estimated using the position of the maximum in the yellow channel. Since this single-pixel value is used in all subsequent calculations, we minimized uncertainty by using the relative shift between the red and yellow channel maxima, based on both buffer images. Each shift yielded an independent estimate of the array position; the final intensity was taken as the average of the values at both estimated locations. The two location estimates typically differed by 1–2 pixels, resulting in certain uncertainty in the kinase activity measurement (median error < 0.03; 80th percentile < 0.08). See figure below.

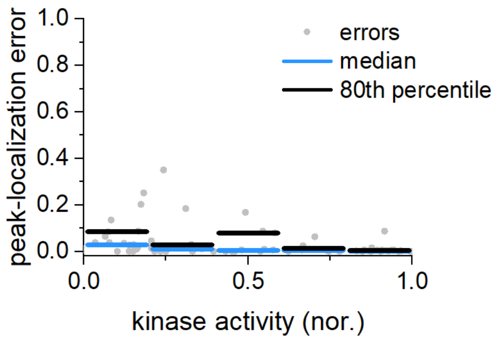


**Single-array dose–response measurements.** The single-array dose-response measurements (Fig. 5D–E) were analyzed as described above. The bleaching factor (α, see Fig. S3) was set for each array such that the red peak in the first and last images (both acquired in buffer) matched. Arrays with α values outside one standard deviation of the mean were omitted from the analysis.

### Supplementary Figures

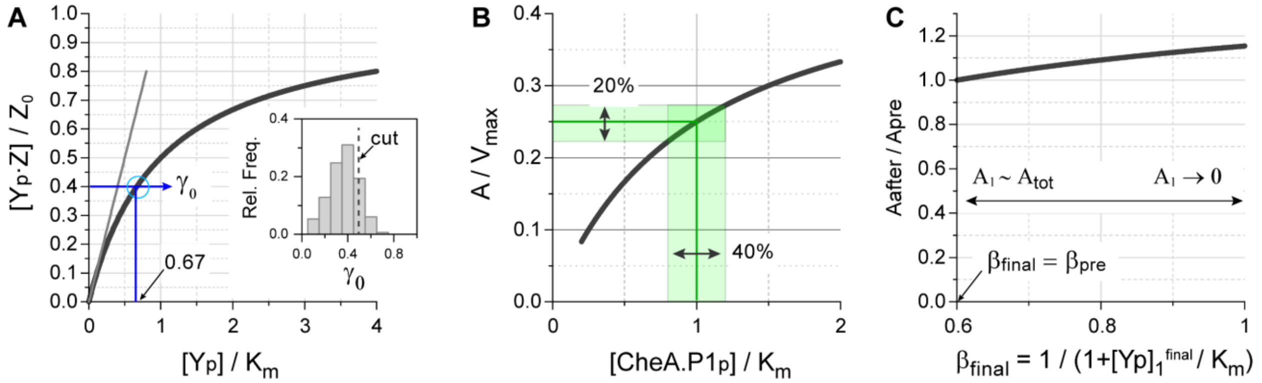


**Figure S1**. Estimating the inherent coupling of kinase activity in different arrays within a single cell (see Supplementary Text 1). (**A**) The fraction of CheZ bound to arrays is plotted against the total CheYp in the cell (black line). The observed distribution of the bound-CheZ fraction in different cells is shown in the inset. A few cells with higher $\gamma_0$ values and more than one dominant array were removed from the analysis (dashed line). (**B**) The fraction of kinase activity is plotted against the resulting CheA·P1 phosphorylation. (**C**) We consider two arrays in the same cell: one remains unstimulated, while the other is fully inhibited by a specific stimulus. The plot shows how turning off the second array ($A_2 \rightarrow 0$) affects the kinase activity in the unstimulated array ($A_1$). The change in $A_1$ is shown as a function of $\beta_{final} \equiv 1/(1 + Yp_{final}/K_m)$, which represents the final level of CheYp in the unstimulated array 1.

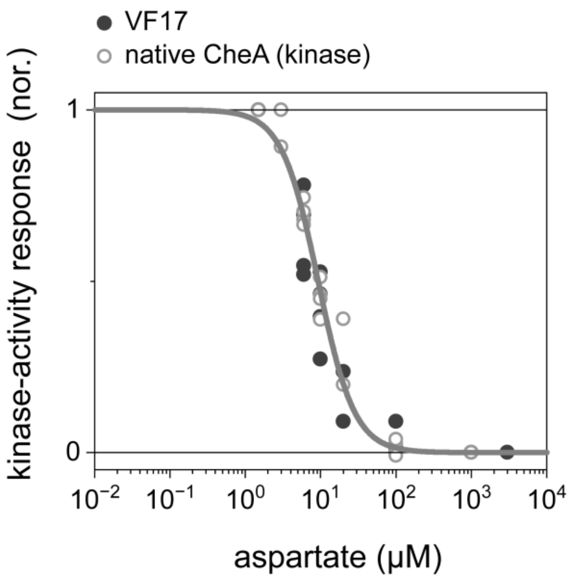


**Figure S2**. Aspartate dose-dependent kinase inhibition measured by population FRET in VF17 cells between CheA::CheY–mYFP and CheZ(F98S)–Scarlet (black symbols), or in V689 cells with native CheA (UU2828, *ΔcheYcheZ*) expressing tagged CheY and CheZ from a plasmid (gray symbols; pAV109). Shown is the change in kinase activity (A) as a function of aspartate concentration: (A([asp])-$A_{sat}$)/(A([0])-$A_{sat}$), where $A_{sat}$ being approximately 22% larger in the VF17 cells, consistent with the shift in $K_{1/2}$ observed in Fig. 1C. These changes correspond to an estimated 3–10% change in Tar fraction, based on equations E2–3, assuming uniformly mixed arrays. The gray line represents a fit to a Hill function with $K_{1/2}$ = 9.2 μM and a Hill coefficient of 1.8.

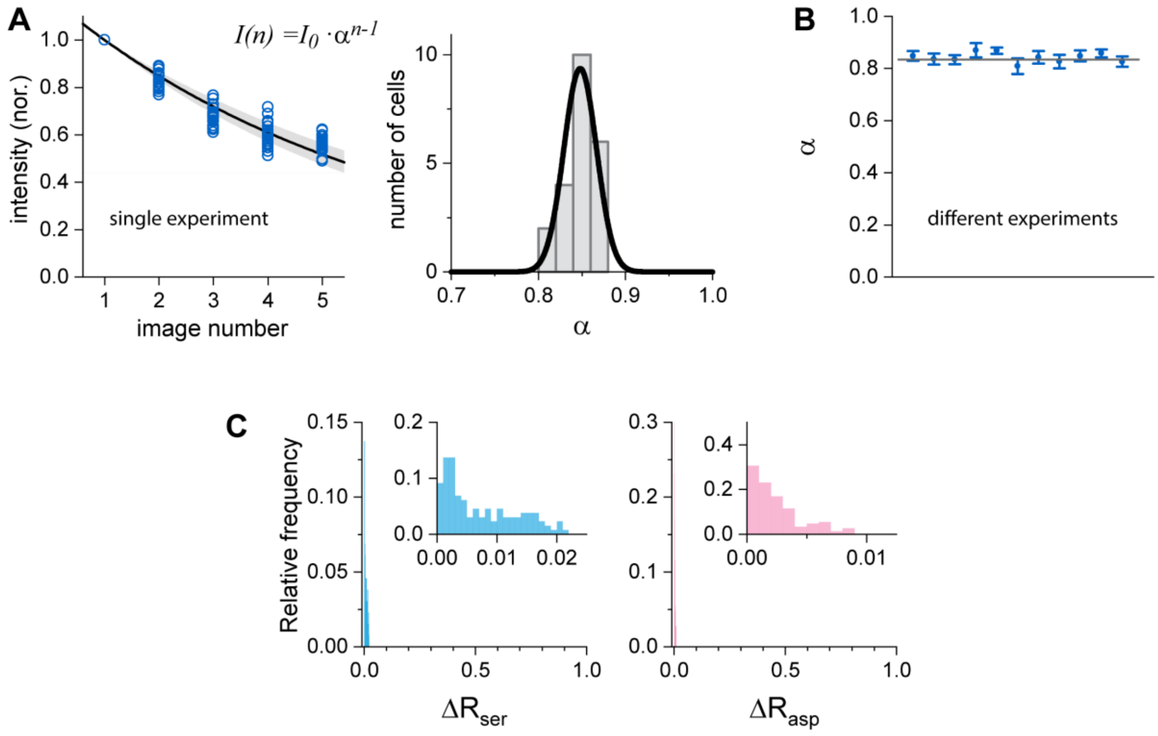


**Figure S3**. Correction for the mScarlet-i bleaching (see Supplementary Text 2). To account for bleaching, we corrected the n-th image according to: $I_{corrected}(n) = I_{meseared}(n)/\alpha^{(n-1)}$. The bleaching factor $\alpha$ was assigned for each experiment based on the population averaged bleaching measurements in the particular experiment. (**A**) Shows the measured intensity for several individual cells (left panel) and the distribution of the corresponding α values in that experiment (right panel). (**B**) The values of $\alpha$ in different experiment. (**C**) Uncertainty in the calculated responses to serine ($R_{ser}$) and aspartate ($R_{asp}$) due to uncertainty in $\alpha$ (see Supplementary Text 2). Shown are histograms of the uncertainties in the responses for a population of 131 clusters. Insets show a zoomed-in view of the same distributions.

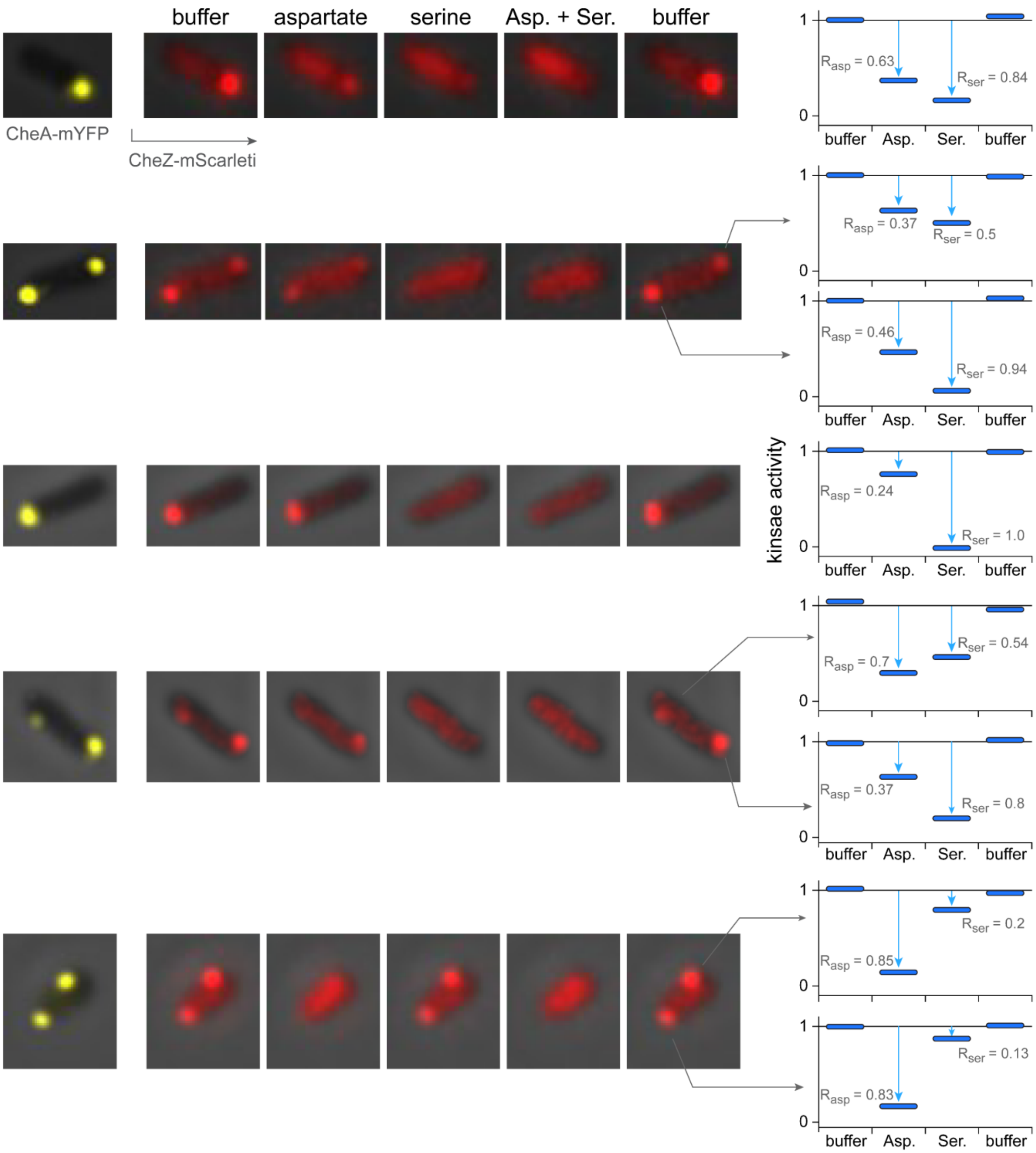


**Figure S4.** Additional examples for single array responses. Images were corrected for photobleaching.

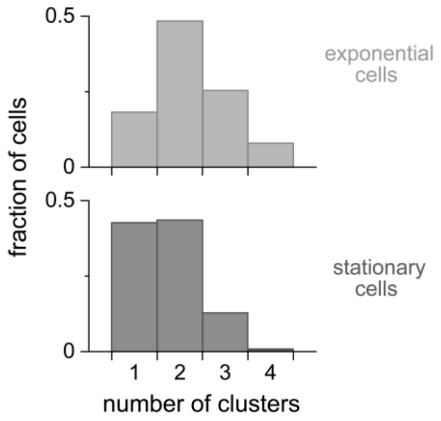


**Figure S5**. Number of arrays identified in each cell. VF17 cells were sampled from exponential-phase culture (OD 0.45; 126 cells) or stationary-phase culture (OD 3.5; 117 cells). Arrays were identified using the mYFP fluorescence images. Note that not all the arrays in each cell were further analyzed for activity responses.

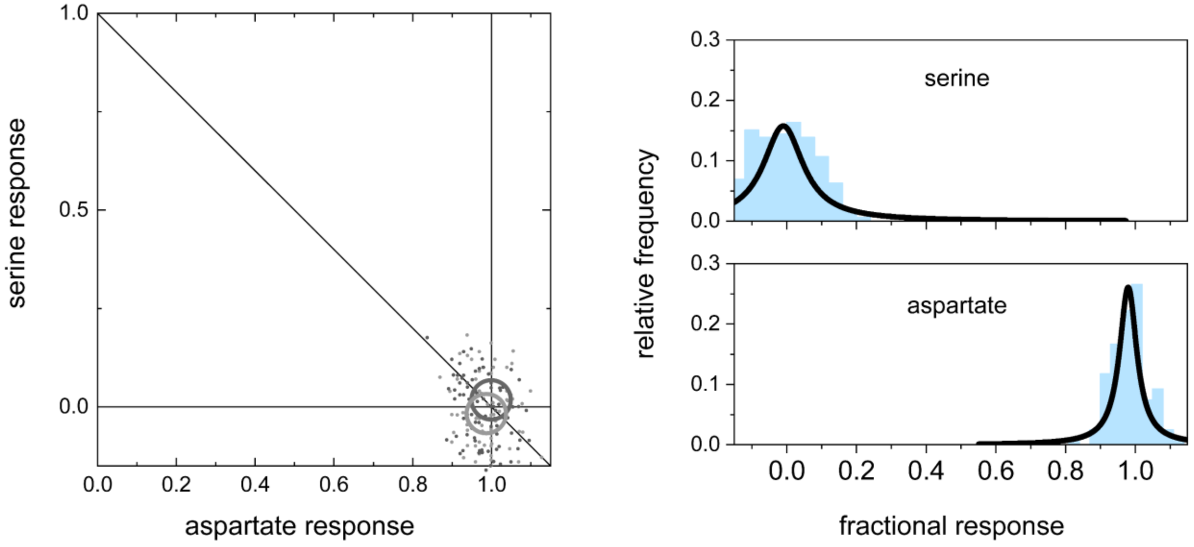


**Figure S6**. Responses of individual arrays in cells lacking Tsr. The response properties of arrays, as in Fig. 2C, measured in MK32 (VF17:Δ*tsr*) cells during the exponential (light gray; OD 0.45) or stationary (dark gray; OD 3.5). The corresponding serine and aspartate response distributions are shown on the right. Circles mark the mean responses and their size represents ±5% deviation.

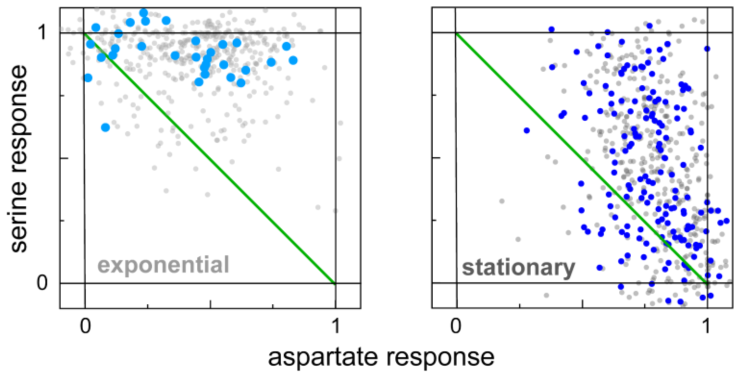


**Figure S7**. Responses of individual dominant arrays. Response properties of arrays as in Fig. 2C, with a sub-population of the arrays highlighted. Highlighted arrays are those that are present alone in their respective cells or are much larger than other arrays within the cell, such that potential CheZ-mediated interference is excluded.

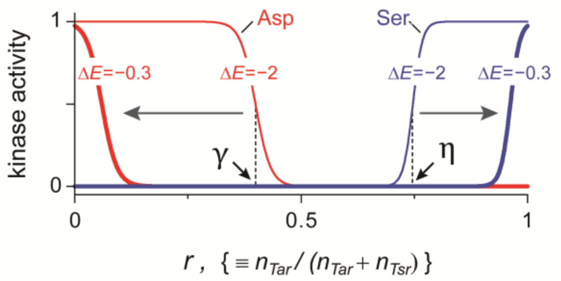


**Figure S8**. MWC-model analysis (see Materials and Methods, Eqs. 2-3). We consider an MWC-cluster containing both Tar ($n_{Tar}$) and Tsr ($n_{Tsr}$) in the presence of saturating aspartate (red lines) or serine (blue lines). The kinase 'on' probability is plotted against the Tar fraction $r \equiv n_{Tar}/(n_{Tar} + n_{Tsr})$, using the following parameters: $n_{Tot} = 12$; $K_{on}/K_{off} = 150$ for Tar and $2{,}700$ for Tsr; and $\Delta E = -2$ (thin lines) or -0.3 (thick lines) for both receptors. Note the shift in the critical points with the energy bias of the receptors (grey arrows).

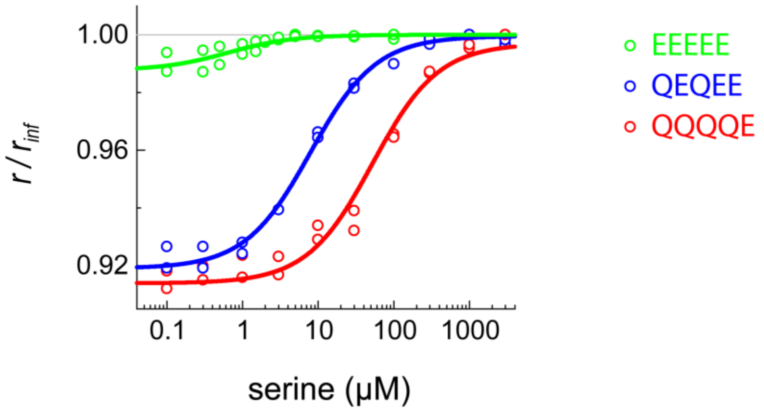


**Figure S9**. Anisotropy dose-response measurements of Tsr to serine. UU1581 cells lacking all chemotaxis proteins (Sandy Parkinson, University of Utah, Salt Lake City) expressing from a plasmid Tsr-mYFP receptors with different E and Q replacements at their modification locations, as labeled. Cells were grown overnight in 1 ml Bacto-Tryptone (TB 10 g/l; NaCl 5 g/l), diluted 100-fold in 10 ml Bacto-Tryptone supplemented with Ampicillin and IPTG, and allowed to grow to $OD_{600}$ ~ 0.45. Cells were washed and resuspended in a motility medium: 10 mM potassium phosphate, 0.1 mM EDTA, 1 μM methionine, and 10 mM lactic acid, pH 7. Following ref. 24, we evaluated the Tsr properties in vivo based on their physical responses to serine, by measuring anisotropy changes, which, as described before, indicate changes in the packing of the receptors, in this case, most-likely within trimers.

The data was fitted (lines) by:

$$P_{on}(L) = \frac{1}{1 + e^{\Delta E} \cdot f(L)} \qquad where \qquad f(L) \equiv \frac{1 + L/K_{off}}{1 + L/K_{on}}$$

And the anisotropy, $r(L)$ was fitted by

$$\frac{r(L)}{r_{inf}} \equiv \frac{r(L)}{r_{off}} = 1 - \left(1 - \frac{r_{on}}{r_{off}}\right) \cdot P_{on}(L)$$

Where L is the serine concentration, $K_{off}$ = 0.55 uM, $K_{on}$ = 1500 uM, $\Delta E_{EEEE}$ = 1.8, $\Delta E_{QEQE}$ = -2.6, and $\Delta E_{QQQQ}$ = -4.58. Experiments were performed at room temperature (22°C).

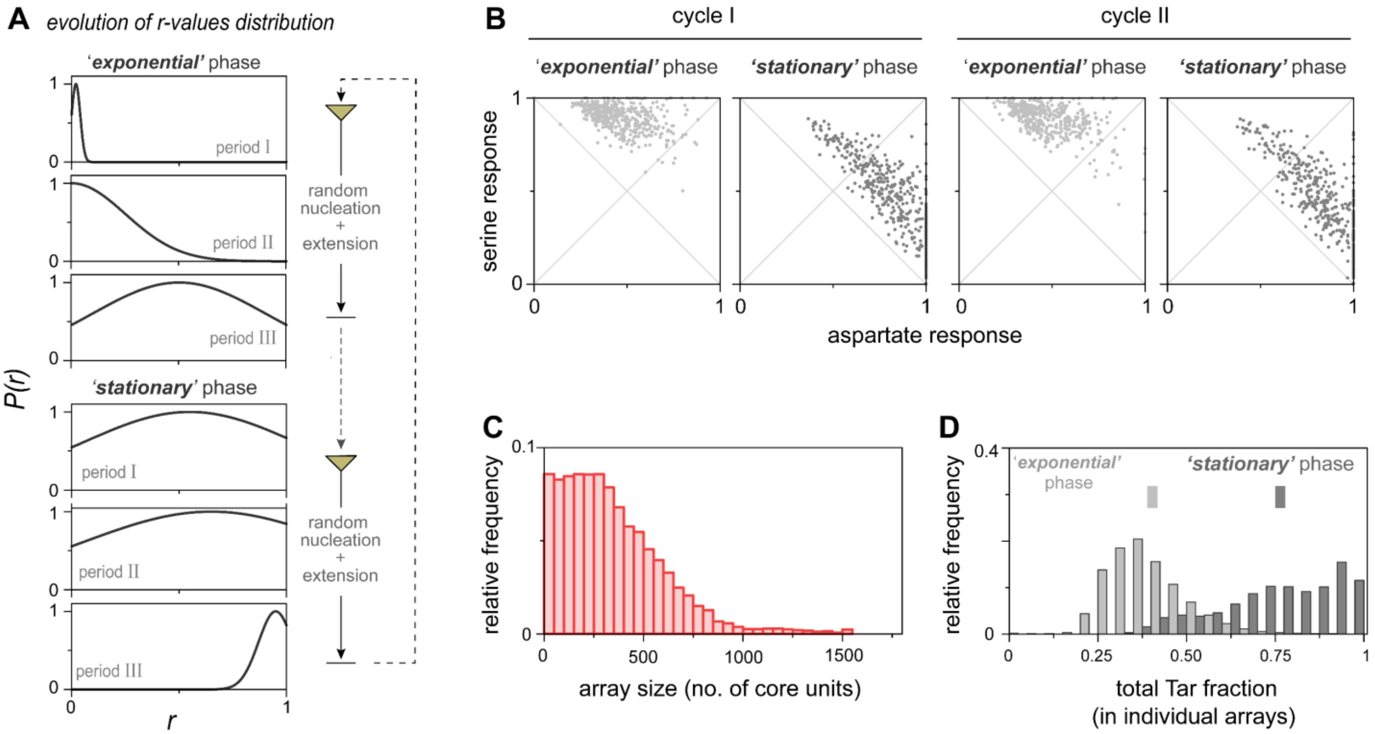


**Figure S10.** Numerically generated arrays with non-uniform Tar and Tsr distributions corresponding to the 'exponential' or 'stationary' growth phases. Arrays were generated as outlined in Materials and Methods (see also Fig. 5). (**A**) A schematic depiction of the procedure, including the *r-value* distributions used in the generation of core unit. In each case, Tar and Tsr were randomly picked with a probability set by the *r-value.* To create temporal correlations, each randomly picked *r-value* was, on average, for the creation of 10 core units. The number of arrays increased 8-fold and 4-fold in 'exponential' and 'stationary' phases. (**B**) The response properties of the arrays were analyzed at the end of each section using Eq. 1 (Materials and Methods), using the following parameters: the ratio $K_{on}/K_{off} = 150$ for Tar and 2,700 for Tsr; and $\Delta E = -0.8$ for Tar and $-2.6$ for Tsr. and the size of the cooperative regions (N) was 4 core units. All energies are in units of $\mathrm{k_B T}$. Two consecutive 'growth' cycles were done. (**C**) The distribution of array sizes. (D) The overall fraction of Tar receptors in each array is shown for the two 'growth' phases. The population averaged *r* values are also marked, showing ~ 2-fold increase in the stationary phase.

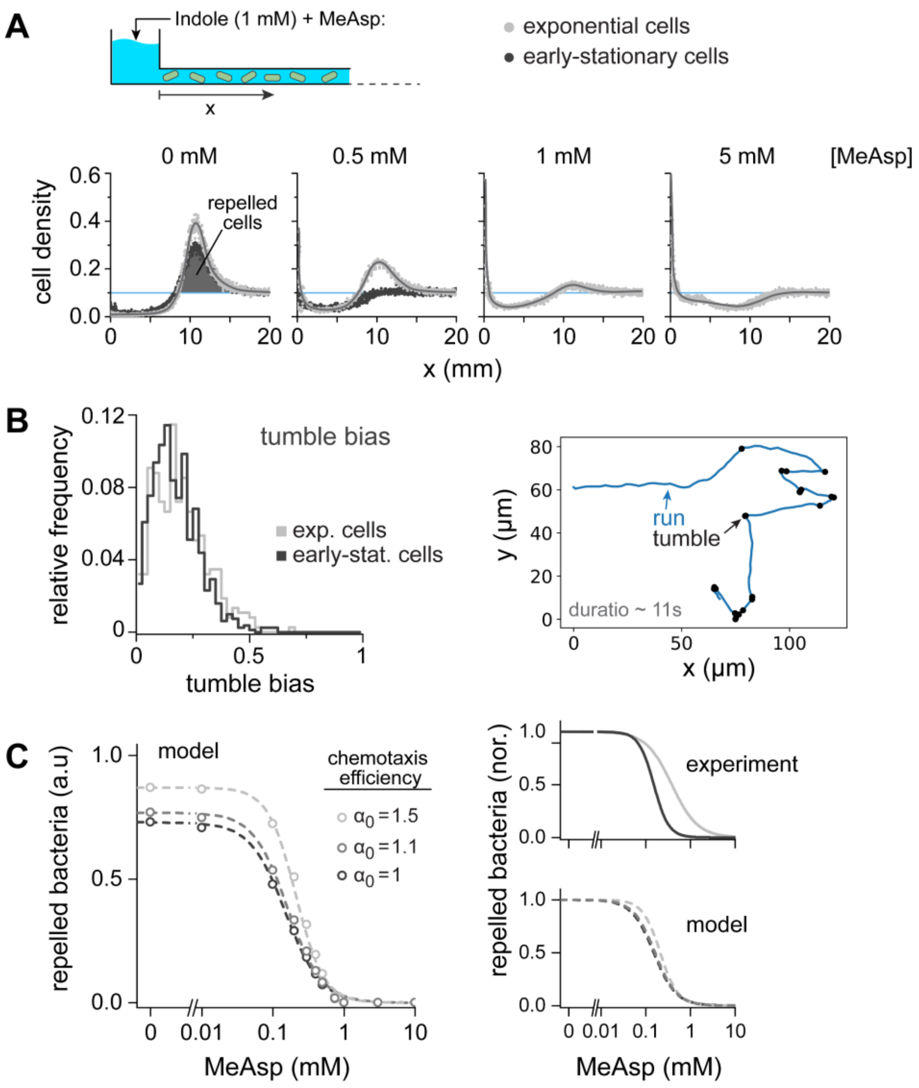


**Figure S11**. Behavioral experiments (see Fig. 6).

(**A**) Representative raw data. Bacterial distribution in the channel 3 hours after introducing the source, for experiments with exponential-phase cells ($OD_{600}$ 0.4; light gray) and early-stationary-phase cells ($OD_{600}$ 1.7; dark gray). Three repetitions are shown for each condition. The source contained Indole (1 mM) and varying concentrations of MeAsp (as labeled). Initially, cells were uniformly distributed at $OD_{600}$ 0.1 (blue line). Lines are a guide to the eye. See Materials and Methods and ref. 40 for detailed information.

(**B**) Distributions of tumbling bias of unstimulated cells in the exponential ($OD_{600}$ 0.4; light gray) and early-stationary ($OD_{600}$ 1.7; dark gray) growth phases (left plot; N = 375 and 822 tracks in exp. and early stat. phases, respectively). Tumbling bias, defined as the fraction of time cells spend tumbling, has been evaluated by single-cell tracking at ~15 frames per second in μ-slide (ibidi 80326), using a 10x (NA 0.45) objective. The mean tumbling bias were 0.195 and 0.174 for the exp. and early-stat. cells (i.e., ~10% shift). A representative track of a single cell is shown on the right with tumbling events marked in black.

(**C**) Since any change in tumble bias is expected to influence both Tsr-mediated and Tar-mediated chemotaxis, it might be expected in this case to have weaker influence on the balance between the responses to the attractant and repellent. To evaluate more quantitatively whether the 10% change in the averaged tumble bias may, by itself, lead to the behavioral change observed in Fig. 6 we employed the mean-field chemotaxis model described in ref. 40: briefly, we numerically solved a 1D chemotaxis model in the case of a constant source emitting two conflicting effectors:

$$\frac{\partial \rho}{\partial t} = D\frac{\partial^2 \rho}{\partial x^2} - \frac{\partial}{\partial x}\left(\rho \frac{\partial}{\partial x}\left(\alpha_0 \cdot \sum_{i=Indole,MeAsp} \alpha_i \cdot \ln\left(\left(k_{off}^i + c_i\right)/\left(k_{on}^i + c_i\right)\right)\right)\right)$$

Where $\rho$ is the cell density, $c_i$ are the ligand concentrations, $D$ is the cells' effective diffusion coefficient, $k_{off}^i, k_{on}^i$ are the effective dissociation constants for the receptors in the ON or OFF states for each ligand, $\alpha_i$ are the chemotaxis coefficients related to each signal, and $\alpha_0$ is set to be a general correction for these chemotaxis coefficients. The model was solved numerically (as in ref. 40) to obtain cell-density profiles in the channel for t = 3h, and the repelled bacteria were evaluated as in Fig. 6C.

The number of repelled bacteria for three values of $\alpha_0$ (1, 1.1, and 1.5) are shown in the main plot (left), and the corresponding normalized plots are shown on the lower-right plot. Also shown in the upper-right plot are the normalized Hill fits to the measured dose-dependent behavior shown Fig. 6C. Clearly, a 10% change in chemotaxis efficiency (i.e., $\alpha_0 = 1.1$) only slightly effecting the behavior and almost no effect after normalization.

Moreover, even if a 10% change in tumble bias were to cause a much larger effect on chemotaxis efficiency (e.g., 50% change; corresponding to $\alpha_0 = 1.5$), it could not account for the observed behavior (see normalized behaviors). Evidently, even such enhancement of the chemotaxis coefficient may only account for the enhanced repulsion in the absence of MeAsp (left plot), but it cannot account for the observed MeAsp dependence (right plots): the $K_{1/2}^{MeAsp}$ shifted from 150 to 380 μM in the experiments (250%) but only from 150 to 210 μM (140%) in the model, and the slope of the MeAsp response (Hill coefficient) clearly decreased in the experiment (from 2.6 to 1.5) but increased in the model (from 1.75 to 2.2).

Overall, under the conditions tested here, the tumble bias distribution did not change much between exponential (OD ~ 0.4) and early exponential (OD ~ 1.7) and it is unlikely to account for the observed behavioral shift that can more naturally link to the observed shift in the sensory properties of the cells.

<u>Parameters used</u>:

$D = 2.5 \cdot 10^{-6}\ cm^2s^{-1}$, $\alpha_{Indole} = 16 \cdot 10^{-6}\ cm^2s^{-1}$ , $k_{off} = 1\ mM, k_{on} = 20\ \mu M$, $\alpha_{MeAsp} = 20 \cdot 10^{-6}\ cm^2s^{-1}$ , $k_{off} = 20\mu M, k_{on} = 20mM$ (21). Ligand concentrations $c_i(x,t)$ were estimated by $c_i(x,t) = c_i^0 \cdot erfc(x/\sqrt{4D_it})$ with $c_{Indole}^0 = 1\ mM$ , $D_{Indole} = 8 \cdot 10^{-6}\ cm^2s^{-1}$ and $D_{MeAsp} = 7.5 \cdot 10^{-6}\ cm^2s^{-1}$.
